# Interspecies interactions drive bacterial proteome reorganisation and emergent metabolism

**DOI:** 10.1101/2025.05.14.653412

**Authors:** Stephan Kamrad, Simone Mozzachiodi, Sonja Blasche, Simran K. Aulakh, David Scheidweiler, Arianna Basile, Rui Guan, Rob Bradley, Naomi Iris van den Berg, Michael Mülleder, Markus Ralser, Kiran R. Patil

## Abstract

Species in microbial communities must respond to the presence of others to stave off resource competition or to capitalise on new resources that may become available due to additional metabolic activities. Such metabolic interactions manifest in growth effects such as competitive exclusion or cooperative growth boost. However, intra-cellular molecular changes that underpin these responses are very sparsely studied, preventing mechanistic insights into community function and dynamics. Here, we analyse meta-proteomics and metabolomic responses in 104 pairwise co-cultures of 15 diverse gut bacteria in a nutrient-rich medium. These co-cultures span a diversity of ecological interactions with competition, amensalism and exploitation being prominent, and with around one in five positive interactions. The species in co-cultures feature substantial molecular responses, with typically more than 60% of the proteome changing in response to at least one partner. Even closely related species and ortholog proteins show different expression profiles in response to the same partner indicating functional diversification at both protein and species level. We show that the magnitude of these responses is determined by genome size, species abundance, and pH. Small-molecule transport and carbon metabolism are among the most responsive processes indicating pervasive metabolic interactions. Using metabolomics, we identify likely cross-fed metabolites, emergent polyamine metabolism, and niche partitioning in amino acid utilisation. Overall, our study shows how bacteria respond to the presence of other species through extensive remodelling of their proteome and metabolome.

## Introduction

Comprehensive catalogues of the composition of diverse microbial communities important for the health and environment are increasingly becoming available [1–6]. An outstanding challenge is to understand mechanisms underlying interspecies interactions that determine the compositional and functional dynamics of these communities in response to biotic and abiotic perturbations [7,8]. However, molecular analysis at community-scale is challenging due to the compositional and dynamical complexity of natural communities. Reductionist (‘bottom up’) approaches, and in particular pairwise interaction analyses [9–15], are therefore instrumental in unravelling mechanisms shaping community-level, emergent functions [16,17].

We here use gut bacterial isolates to probe the molecular basis of interspecies interactions. Ecological interactions among gut bacteria comprise complex competitive and cooperative interactions across several trophic levels [18–22]. For example, many species collectively contribute to the biosynthesis of short chain fatty acids (SCFA) [23,24] and tryptophan-derived metabolites [25]. Gut bacteria therefore present an attractive model to study how community outputs emerge from the interactions of microbial species and their metabolic activities.

A key question is how enzyme abundances and metabolite production are regulated within the community context. Only a few studies [26–29] have systematically investigated functional interactions at the level of gene expression, revealing which subset of the genetic (e.g. enzymatic) repertoire is realised in specific conditions [30]. In this context, proteomic measurements are more accurate readouts of functional state than transcriptomics as they capture post-transcriptional regulation and provide direct measurements of the molecules that carry out the function, e.g. a metabolic reaction. Recent advances in proteomics precision, depth and throughput [31,32] enable the application of proteomics to species mixtures [33–39]. Previous, small scale studies investigating the regulation of gene expression in synthetic gut bacterial communities *in vitro* [40] or in mouse [41,42] revealed substantial proteome remodelling and niche specialisation depending on community context. This is further supported by a microarray-based meta-transcriptomics study of human stool samples [43]. Overall, while the importance of species interactions in shaping microbial gene expression and metabolism are widely acknowledged, there is a striking lack of large-scale studies, limiting the current understanding of molecular processes underpinning interspecies interactions.

We here employ systematic meta-proteomics and metabolomics of binary co-cultures to map physiological changes that emerge between 104 pairs of 15 diverse human gut bacteria. A substantial fraction of proteins responds to the presence of at least one other species, in particular transporters, metabolic enzymes as well as many un-annotated/understudied proteins. We complement this data with metabolomics to identify putative exchanged and emergent metabolites. Our systematic approach demonstrates a framework and resource to uncover the mechanistic basis of interspecies interactions.

## Results

### Combinatorial co-cultures for the systematic molecular characterisation of species interactions

To study the functional adaptation of bacterial gene expression and metabolism in response to other community members, we selected 15 human gut bacteria (**Fig 1A**, <u>Supplementary Data 1</u>). These include abundant and prevalent commensal species from four phyla, as well as two common pathogens, *Clostridioides difficile* and *Klebsiella aerogenes*, and two probiotics, *Lactobacillus acidophilus* and *Lactobacillus gasseri.* Strains were cultured in a pairwise combinatorial design in 96-well plates, resulting in 608 samples (after quality control) across 104 species pairs (n`≈`4 biological replicates, plus n`≈`16 biological replicates of each mono-culture as control). At stationary growth phase, optical densities at 600 nm (OD) were determined and cell pellets and supernatants were harvested and processed for proteomic and metabolomic measurements by liquid chromatography tandem mass spectrometry (LC-MS^2^).

**Figure 1:**
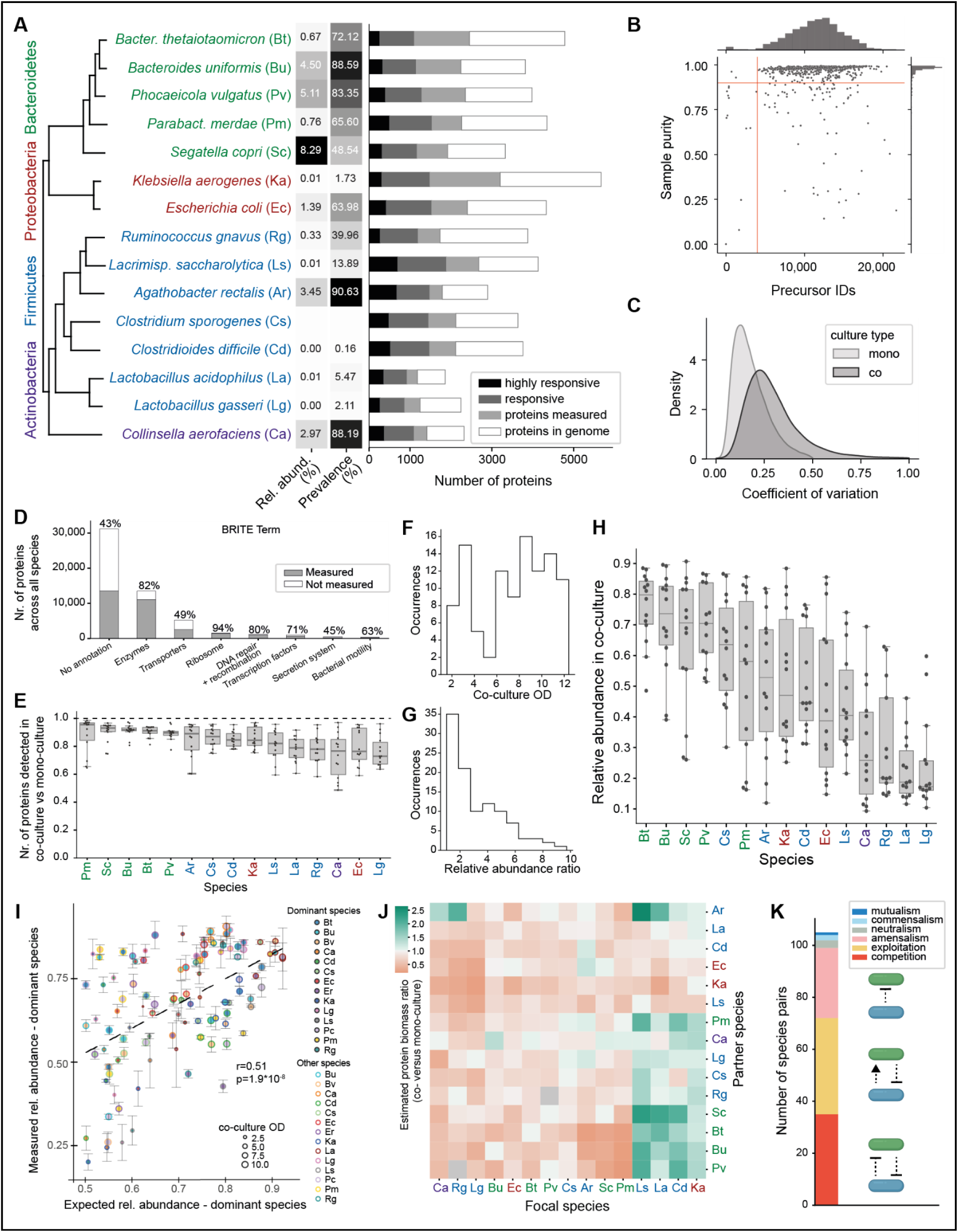
Extensive regulation of proteomes in pairwise co-cultures of human gut bacteria. (A) 15 bacterial strains were selected based on abundance, prevalence and genetic diversity. Proteomes of all pairwise co-cultures were obtained by DIA-proteomics, The phylogenetic tree was generated with OrthoFinder using protein sequences. Bacterial phyla are highlighted in colours. A two-letter code for each species used throughout this manuscript is indicated in parentheses. Mean relative abundance and prevalence are shown in the heatmap and were obtained from a curated set of metagenomic studies [6]. The bar chart illustrates the number of proteins encoded in each genome, the proteins measured in our dataset, the responsive proteins (proteins which are differentially abundant in at least one co-culture, p_adj_<0.05, abs(log_2_(FC))>0.5) and highly responsive proteins (differentially abundant in at least 4 conditions). (B) Quality control of proteomics samples. 768 samples were acquired using dia-PASEF and searched against a combined predicted spectral library of all species. The median number of identified precursors was 17,625. Sample purity was estimated as the fraction of identified precursors assigned to the species expected in the sample. 608 samples with >4,000 identified precursors and a purity of >90% were taken forward. (C) Dataset-wide distributions of CV values across mono-cultures (median=16%, reflecting biological/technical noise) versus co-culture conditions (median=27%, indicating substantial additional variation in response to co-culture partners). (D) Coverage of different functional protein classes (BRITE) by our dataset which includes 31,127 proteins across all species. (E) Fraction of proteins detected in co-versus mono-culture controls. Good data completeness is maintained across co-cultures despite increasing sample complexity. (F) Distribution of total co-culture biomass in OD_600_ units at time of cell collection (24 h). (G) Distribution of relative abundance ratios (major species / minor species) across co-cultures. Relative abundance was estimated from peptide intensities (Methods). (H) Relative abundance of each species across 14 co-culture conditions. Each point represents the mean of n≈4 biological replicates. Some species were consistently more abundant than others, although no single species was always or never the major partner. Box plot elements are defined as usual: center line, median; box limits, upper and lower quartiles; whiskers, 1.5x interquartile range; points, outliers. (I) Relative abundance in co-cultures predicted using ratio of mono-culture OD_600_ correlates poorly with observed relative abundance across all species pairs. I.e. species with similar mono-culture OD_600_ values can form co-cultures with widely varying ratios indicating prevalence of interactions. r = Pearson correlation coefficient. (J) Inter-species growth interactions (absolute protein biomass in co-/mono-culture, log_2_-transformed) of partner species (y-axis) on focal species (x-axis). Negative interactions dominate, but a small group of species (*L. saccharolytica, L. acidophilus, C. difficile, K. aerogenes*) benefits from the presence of certain Bacteroidetes and *A. rectalis*. Grey indicates data not available. (K) Types of ecological interactions found across 104 species pairs. A mean change in growth of >10% was considered a positive/negative effect. Competition, exploitation and amensalism are the predominant forms of ecological interactions across the dataset. Icons are by BioRender.com.

To determine cellular proteomes, we used a throughput-optimised workflow on an Evosep One chromatography system with data-independent acquisition by parallel accumulation serial fragmentation (dia-PASEF [44]) on a trapped ion mobility spectrometry - time of flight (timsTOF) mass spectrometer. Data were analysed with DIA-NN [45], using a concatenated library-free approach, which successfully deconvoluted the two-species proteomes (Methods, <u>Supplementary Data 2</u>). Out of 768 samples for which data was acquired, 106 samples were excluded because they had fewer than 4000 precursor identifications or more than 10% of identified precursors were originating from species other than those expected in the sample (**Fig 1B**). In mono-cultures, after quality filtering (Methods, **SuppFig 1A**), we quantified over 2,000 proteins (56% of the proteins encoded in the genome) per species on average (**Fig 1A**). The median, protein-level coefficient of variation was 16% across mono-culture controls and 27% across co-cultures, indicating low experimental noise levels and substantial biological responses in co-culture (**Fig 1C**). The number of identified proteins compares favourably with other current, state-of-the art studies in bacterial model species [46,47] and indicates high proteome coverage despite short analysis times. Our dataset, like other proteomic datasets, covers the vast majority of the proteome by mass, since highly abundant proteins are much more likely to be measured. E.g., Schmidt et al [48] estimated that the 55% of *Escherichia coli* ORFs detected in their dataset cover 95% of the proteome by mass.

At the functional level, we obtained excellent coverage of annotated enzymes, ribosomal and DNA replication/repair proteins (>80%), good coverage of annotated transcription factor and motility proteins (>70%), but comparatively sparser coverage of membrane proteins (transporters and secretion systems) as well as unannotated proteins (**Fig 1D**). In co-cultures we typically detected a much higher number of proteins overall, but the coverage per individual species declined slightly to 77%-92% of the mono-culture proteins (first and third quartile, **Fig 1E**). This is expected as the abundance of peptides from each species is lower and the sample complexity is higher for two-species meta-proteomes in comparison to single-species proteomes. In summary, we collected a systematic, broad and deep meta-proteomics dataset of bacterial co-cultures.

### Diverse ecological interactions emerge in co-cultures

First, we leveraged the proteomics data and co-culture ODs to determine abundances of individual species in co-culture. OD values of co-cultures, representing total biomass, typically ranged from 3.1 to 9.7 (first and third quartile, respectively) and were somewhat uniformly distributed (**Fig 1F**). For all co-cultures, we computed the absolute abundances of each member species based on species-specific peptide intensities and the OD of the co-culture (Methods, see also [28]). In the majority of co-cultures (64%), one species was substantially more abundant than the other (>2x), although in only a small minority (19%) the ratio was more than 5 (**Fig 1G**). While no single species was always or never the dominant partner across all co-cultures, a few species were usually the major species (the Bacteroidetes *Phocaeicola vulgatus, Bacteroides uniformis, Bacteroides thetaiotaomicron and Segatella copri*), and a few others were usually the minor species (the lactobacilli *L. gasseri* and *L. acidophilus*, as well as *Collinsella aerofaciens* and *Ruminococcus gnavus*) (**Fig 1H**). The growth of a species in mono-culture (final OD) was able to explain some of the relative abundance in co-culture (r=0.52) but for species with similar OD in mono-culture the prediction was poor (**Fig 1I**), indicating prevalent inter-species interactions.

The majority (134/223) of growth interactions across all co-cultures were negative (>10% decreased absolute abundance in co-versus mono-culture) consistent with the resource limitation in batch cultures, yet a significant minority (41/223) was positive (>10% increase in growth) and the rest (48/223) neutral (**Fig 1J**, **SuppFig 1EF**, <u>Supplementary Data 3</u>). We identified four phylogenetically diverse species, *Lacrimispora saccharolytica, C. difficile, L. acidophilus* and *K. aerogenes*, which benefit from the presence of Bacteroidetes and *Agathobacter rectalis*. The magnitude of these interactions is substantial; e.g., *L. saccharolytica*, which had a biomass of 1.38±0.06 OD units in mono-culture, grew to 3.66±0.21 in the presence of *A. rectalis* and to 3.21±0.22 in the presence of *S. copri*. We independently validated the growth promotion of *C. difficile* by four Bacteroidetes species using quantitative flow cytometry (**SuppFig 2**).

**Figure 2:**
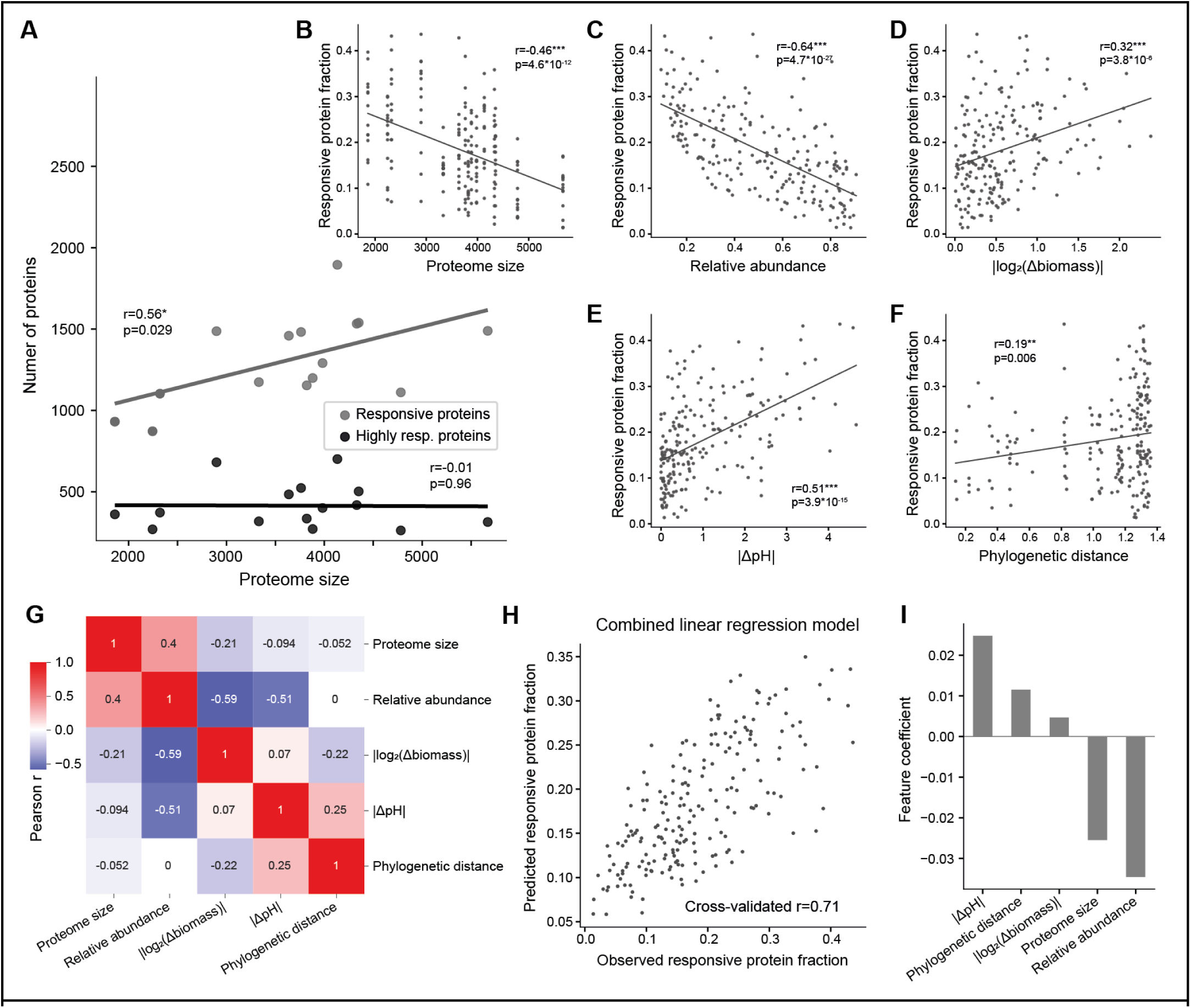
Broad factors underlying magnitude of proteome response. (A) Number of responsive and highly responsive proteins (hit in at least 4 co-culture conditions) dependent on proteome size at the species level. The number of responsive proteins increases with genome size, although not at the same rate. The number of highly responsive proteins is not correlated with genome size. (B) Responsive proteome fraction (number of hits / number of proteins in proteome, y-axis) across all 208 datapoints (104 co-cultures, from the perspective of each species) versus proteome size (number of proteins encoded in genome). (C) Same for relative abundance (fraction of protein biomass in co-culture attributable to species). (D) Same for | log_2_(Δbiomass) | - The absolute log_2_-transformed fold change of the biomass reached in co- versus mono-culture, reflecting ecological interactions with the partner species. (E) Same for | ΔpH | - The absolute change in pH between the co-culture and the mono-culture. (F) Same for the phylogenetic distance between the co-culture partners, obtained from the tree shown in **Fig 1A**. (G) Heatmap indicating Pearson correlations between the factors shown in C-F. (H) A multiple linear regression model based on the factors from C-F was used to predict the responsive proteome fraction. Leave-one-out cross-validation was used to assess the ability of the model to predict unseen interactions. (I) Model coefficients indicate the importance of each shown feature in predicting the magnitude of the proteome response. The proteome size of the focal species and changes in pH were identified as major drivers, followed by relative abundance. Changes in growth and phylogenetic distance between the partner species on the other hand had a minor impact. For A-F,H: The Pearson correlation coefficient is indicated in the plot, fitted lines were obtained by ordinary least squares regression. Significance keys: ***p<0.001, **p<0.01, *<p<0.05

Across all the co-cultures, 33% of pairs could be classified as competitive (absolute abundance of both species lower than in their respective mono-cultures) and 27% as amensal (one species inhibited, the other unchanged) (**Fig 1K, SuppFig 1E**). However, strikingly, the most common (35%) interaction type was exploitation (where one species increased in abundance, the other decreased). Together with two commensal and one mutualistic species pairing, 37% of pairings involve a positive outcome for a member species. Overall, our co-culture model produced a broad diversity of growth behaviours and ecological interactions.

### Extensive proteome remodelling in the co-culture

To investigate the molecular changes associated with the various observed ecological interactions, we identified proteins that were differentially abundant in co-cultures, in comparison to the corresponding mono-cultures (log_2_-transformed absolute fold change >0.5, FDR-adjusted p-value <0.05, moderated t-test via *limma*, **SuppFig 1B-D**, <u>Supplementary Data 2</u>). In 1163 instances (0.25% of the dataset), a protein was only identified in co-culture but not in mono-culture (indicating a likely upregulation of these proteins in co-culture) and these were also classified as hits.

Between 45% (*B. thetaiotaomicron*) and 83.1% (*A. rectalis)* of measured proteins (median=68.8%) were differentially expressed in at least one co-culture (**Fig 1A)**. Overall, the proteome response was highly specific to the co-culture partner (33% of responsive proteins only differentially abundant in a single co-culture), however, we also identified common responsive proteins (hits in 4 or more conditions), which made up between 9% (*K. aerogenes*) and 38% (*A. rectalis*) of measured proteins (median 21%).

### Physiological factors underlying strength of proteome response in co-cultures

In the search of a mechanistic basis and general principles underlying the proteome changes observed in co-culture, we investigated a range of factors ranging from intrinsic properties of a species like its proteome size and extrinsic factors like pH change (Supplementary Data 4).

**(i) Proteome size** - The number of proteins encoded in the genomes of the selected species varies more than two-fold; between 5670 for *K. aerogenes* and 1859 for *L. acidophilus* (median=3820) (**Fig 1A**). Proteome size was correlated with the total number of responsive proteins across co-cultures but not the number of highly responsive proteins (**Fig 2A**). Thus, across diverse species, there is a consistently large set of approximately 400 proteins that is highly regulated in response to other species. As proteome size increases, the number of regulated proteins increases at a lower rate, resulting in a negative correlation between proteome size and the responsive protein fraction (the number of hits divided by the number of quantified proteins (**Fig 2B**). **(iii) Relative abundance** – We note an appreciable correlation between relative abundance and the fraction of differentially expressed proteins (**Fig 2C**), potentially because relatively low abundant species are more exposed to the other species’ cells, secreted proteins and metabolites than their own.
**(ii) Growth** - Changes in growth are known to be linked with proteome changes [48,49]. Indeed, the responsive protein fraction correlates with the magnitude of change in biomass of the species in co-culture compared to mono-culture (**Fig 2D**). However, even low abundant species can have a pronounced effect on the protein expression of high abundant species. For example, the dataset-wide strongest response was observed in *A. rectalis* when co-cultured with *E. coli* (21.8% of proteome and 37.7% of measured proteins responding), although *A. rectalis* was the strongly dominant species (relative abundance of 80.5%). **(iv) pH** - Bacterial metabolism commonly produces overflow metabolites like short chain fatty acids and other organic acids that alter the pH of the surrounding environment. We assessed the pH of the co-cultures at the time of proteomic and metabolomic sampling using an indicator dye (Methods). In monocultures, we observed strong media acidification by Bacteroidetes and *A. rectalis* (**SuppFig 3**), compatible with their known capacity for producing organic acids during carbohydrate fermentation [50,51]. The *Clostridia C. sporogenes*, and *C. difficile* increased the pH, attributable to ammonia production during Stickland metabolism of amino acids [52,53]. *L. saccharolytica* and *C. aerofaciens* likewise raised the pH. In co-cultures, strong changes in pH compared to the mono-culture were associated with an increased number in differentially expressed proteins (**Fig 2E**). **(v) Phylogenetics** - Finally, phylogenetic distance between co-culture partners was weakly correlated with the responsive proteome fraction (**Fig 1F**).

To gauge the combined influence that these factors hold, we constructed a multiple linear regression model and performed leave-one-out cross-validation (**Fig 2G**). This model achieved a Pearson r of 0.71 in predicting the responsive protein fraction (**Fig 2H**). Analysis of the model coefficients revealed changes in pH and relative abundance as the most important drivers, followed by proteome size and phylogenetic distance (**Fig 2I**). Thus, both intrinsic and extrinsic factors determine the proteomic response of a species to their co-culture partners.

### Carbohydrate metabolism and transport are highly responsive in co-culture

To identify biological processes differentially regulated in co- versus mono-culture, we used a sequence-based search to map proteins to KEGG orthologous groups and retrieve associated functional annotation (Methods). Globally, the set of responsive proteins (hit in at least one species-condition pair) was significantly enriched in 7 KEGG pathways (**Fig 3A**) (p_adj_<0.05, FDR-corrected Fisher’s exact test). These included specific classes of nutrient transport machinery, namely the phosphotransferase system (PTS) and ATP-binding cassette transporters (ABC). Three other pathways involved in carbohydrate metabolism (butyrate metabolism, starch and sucrose degradation and ‘other glycan degradation’) were also globally enriched, indicating a central role for carbohydrate uptake and metabolism in mediating inter-species proteomic interactions. The set of responsive proteins were depleted in cellular housekeeping functions such as aminoacyl tRNA synthesis, RNA polymerase, homologous recombination and nucleotide metabolism, indicating that these core functions are buffered against communal perturbations.

**Figure 3.**
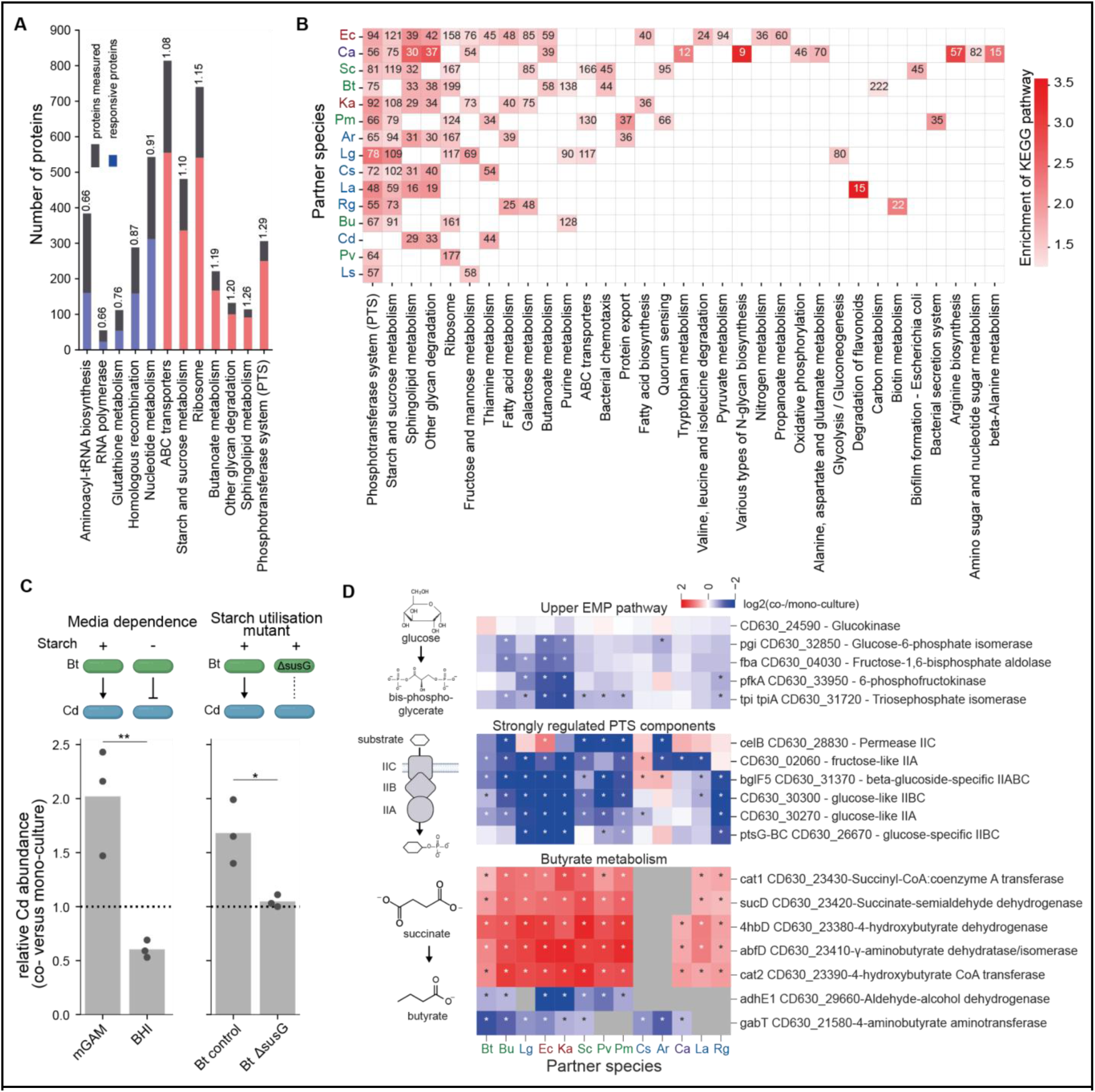
Functional analysis of proteome changes. (A) Global enrichment analysis of KEGG pathways in responsive proteins (differentially expressed in at least one co-culture condition) versus all measured proteins as background. Numbers above the bars indicate the enrichment factor (term frequency in responsive proteins divided by background frequency). Only significantly overrepresented (red) and underrepresented (blue) terms are shown (p_adj_<0.05, FDR-corrected Fisher’s Exact test). (B) Enrichment analysis of KEGG pathways in response to different partner species. Only significantly overrepresented terms are shown (p_adj_<0.05, FDR-corrected Fisher’s Exact test). (C) The bar plot illustrates the stationary-phase abundance of *C. difficile* in co-culture with *B. thetaiotaomicron* divided by *C. difficile* abundance in mono-culture in the indicated conditions, captured by quantitative flow cytometry (**SuppFig 2**). In mGAM, the main medium used in this study, *B. thetaiotaomicron* promotes the growth of *C. difficile* (see also **Fig 1J**) but in Brain Heart Infusion (BHI) medium the relative growth of *C. difficile* is significantly reduced (p=0.0081). A major difference between BHI and mGAM media is the absence of starch. This was further validated by the use of a *B. thetaiotaomicron* starch utilisation mutant (ΔsusG, see Methods) which similarly reduces relative *C. difficile* abundance (p=0.022) in mGAM and appears unable to promote the growth of *C. difficile*. Significance keys: *p<0.05, **p<0.01, Student’s two-sided t-test. (D) Regulation of selected metabolic pathways in *C. difficile* across co-culture conditions. Top: upper Embden–Meyerhof–Parnas/glycolysis proteins; middle: strongly regulated (sum(abs(log_2_(fold-change)))>10) phosphotransferase system components. The vast majority of these are annotated as glucose-specific (see **SuppFig 4C** for other PTS components); bottom: enzymes of the pathway that converts succinate to butyrate.

Zooming in, we next asked which species trigger differential expression of which pathways in their co-culture partner (**Fig 3B**). Many species induced changes in the PTS system (**SuppFig 4A+C)**, ABC transporters (**SuppFig 4B**), starch and sucrose metabolism, sphingolipid metabolism and ‘other glycan degradation’. *L. saccharolytica* had an outstandingly high number of (regulated) ABC transporters. Polysaccharide utilising loci (PULs) on the other hand were not strongly regulated in *B. thetaiotaomicron*, with only three loci out of 23 showing strong regulation across conditions (**SuppFig 4D**). Other pathways were regulated in response to particular species. E.g., arginine biosynthesis was often regulated in the presence of *C. aerofaciens* (**SuppFig 4E**) which depletes arginine from the media (see below). Thiamine biosynthesis was regulated when *C. sporogenes*, known to display thiaminase activity [54], was present in the co-culture (**SuppFig 4F)**. Overall, our data uncovered carbohydrate metabolism and nutrient transport pathways that are extensively regulated in response to the presence of other species.

As a major pathogen, ecological interactions of *C. difficile* with other gut bacteria have been studied intensely [14,55–57]. In one recent study [58], *Bacteroides* and *Phocaeicola* were identified as suppressors of *C. difficile* when cultured continuously in Brain Heart Infusion (BHI) with additional supplements, attributed partially to competition for amino acids. We were thus surprised to see a growth-promoting effect of several Bacteroidetes and *A. rectalis* on *C. difficile* (**Fig 1J**). A major difference between the studies is the cultivation medium, and indeed we observed that *B. thetaiotaomicron* became growth-inhibiting instead of -promoting in BHI (**Fig 3C**). Since mGAM contains substantial amounts (12 g/L) of starch, we tested if starch metabolism by *B. thetaiotaomicron* affects *C. difficile* growth in co-culture by constructing a mutant lacking a key component of the starch utilisation system (*sus)*, the α-amylase susG (Methods, **SuppFig 5**). This mutant did not promote the growth of *C. difficile* (**Fig 3C**), clearly implicating starch metabolism by *B. thetaiotaomicro*n in this interaction.

What molecular cross-feeding interactions drive this interaction? Starch breakdown by sus proteins initially proceeds extracellularly [59], potentially releasing common goods. The proteomics data indicates down-regulation of PTS sugar transporters and upper Embden–Meyerhof–Parnas/glycolysis proteins in the presence of many partner species and strong upregulation in enzymes involved in butyrate production (**Fig 3D**), consistent with previously described cross-feeding of succinate [60] and acetate [18] from *Bacteroides* to *Clostridia*. Related to this, the pH of *C. difficile*- *Bacteroides* co-cultures is acidic, potentially alleviating self-inhibiting effects of the high pH observed in *Clostridium* mono-cultures. Overall, this illustrates the value of proteomic data for generating hypotheses about exchanged metabolites and we next set out to further investigate interactions at the metabolomic level.

### Targeted metabolomics reveals emergent metabolism and cross-fed metabolites

To further the insights into co-culture metabolism, we applied a panel of targeted LC-MS^2^ metabolomic assays (<u>Supplementary Data 5+6)</u> to the co-culture supernatants, covering amino acids, short- and branched-chain fatty acids, tryptophan-derived metabolites and biogenic amines. Comparing metabolite concentrations to fresh mGAM medium, we identified 74 instances where at least one of the tested metabolites was produced by a species in mono-culture (**Fig 4A**, <u>Supplementary Data 7</u>, Methods). As expected, Bacteroidetes produced large amounts of the fermentation product succinate. Many species produced the SCFA propionic acid, as well as the branched chain fatty acids (BCFA) isobutyric and isovaleric acid from valine and leucine. Additionally, we recapitulated well-known specific metabolic activities, such as indole-3-propionic acid (IPA) produced by *C. sporogenes* [61], polyamines by *E. coli* [62] and histamine by *K. aerogenes* [63]. These data are complementary to previous surveys of supernatant metabolomics in gut bacteria [64] as we have used a different growth medium and cover some additional compounds such as SCFA and BCFA.

**Figure 4:**
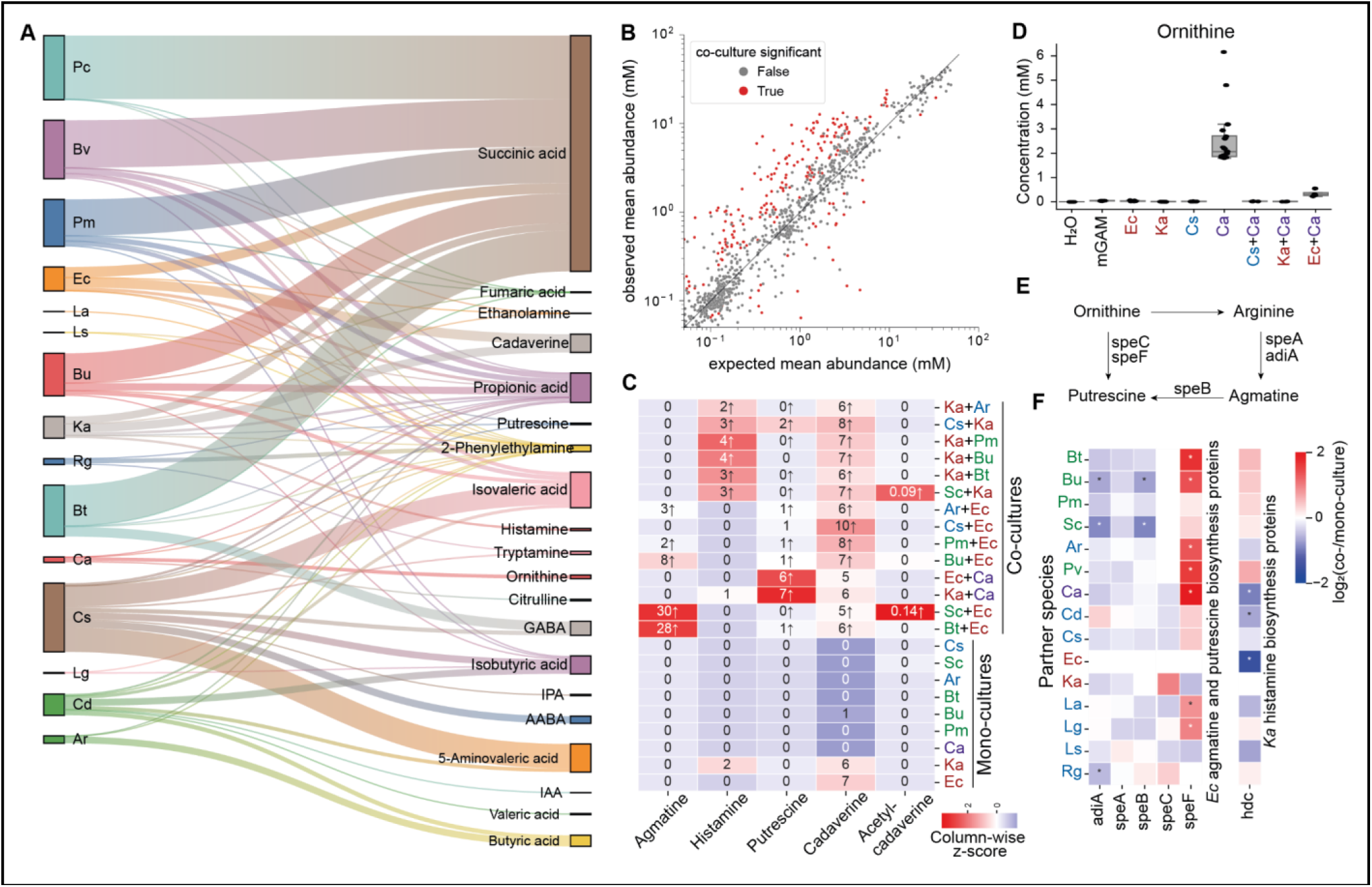
Polyamine and SCFA metabolism are shaped by cross-species interactions. (A) Metabolites produced by individually gut bacteria in mono-culture. Bar width is proportional to concentration. Abbreviations: IPA: indole-propionic acid, IAA: indole-acetic acid, GABA: γ-aminobutyric acid, AABA: α-aminobutyric acid. (B) Comparison of observed and expected metabolite concentrations in co-cultures. Each dot represents a metabolite in a specific co-culture. Expected concentrations were computed from mono-culture concentrations and abundance of species in co-culture. Observed concentrations deviating significantly from expected are shown in orange (p_adj_<0.05 FDR-corrected two-sided Welch’s t-test, log_2_(abs(fold-change))>0.5 and min(observed or predicted concentration)>50 µM or >500 µM for SCFA). (C) Emergent polyamine metabolism in selected co-cultures involving *E. coli* and *K. aerofaciens*. Heatmap colours reflect column-wise z-scores and numbers reflect the mean concentration in µM. *E. coli* does not produce agmatine but does so in co-cultures with specific Bacteroidetes. Histamine production by *K. aerogenes* is stimulated by *C. aerofaciens* and *S. copri*, while putrescine is produced in only co-cultures of *E. coli* or *K. aerogenes* with *C. aerofaciens*. Low levels of acetylated cadaverine emerged in co-culture with *S. copri*. (D) Ornithine accumulates in *C. aerofaciens* mono-cultures but is depleted in co-cultures with *C. sporogenes*, *K. aerogenes* and *E. coli*, possibly indicating cross-feeding. Box plot elements are defined as usual: center line, median; box limits, upper and lower quartiles; whiskers, 1.5x interquartile range; points, outliers. (E) Genes and metabolites involved in agmatine and putrescine synthesis in *E. coli*. Since ornithine can be metabolised to putrescine via ornithine decarboxylases, ornithine is produced by *C. aerofaciens* but depleted in co-culture with *E. coli*/*K. aerogenes*, it is highly likely that cross-fed ornithine is further metabolised to putrescine in these cases. (F) Abundances of enzymes involved in amine synthesis cannot explain the detected concentrations of agmatine, putrescine and histidine. Asterisks indicate hits.

Next, we compared metabolite abundances in co-cultures to expected values based on mono-culture concentrations and relative species abundances under an additive model (**Fig 4B**) (Methods). We identified 267 out of 2600 instances where a metabolite was differentially abundant in a given species pair (p_adj_<0.05 FDR-corrected two-sided Welch’s t-test, log_2_(abs(fold-change))>0.5 and min(observed or expected concentration)>50 µM). In 73 cases, the observed concentrations were below expected, which indicates either competition for precursor metabolites or the partner species consuming the produced metabolite (cross-feeding). In the remaining 194 cases, concentrations were higher than expected indicating upregulation of metabolic activity due to species-species interaction, resource partitioning among co-culture members, or emergent metabolism.

Among the emergent metabolites is GABA, a neurotransmitter and key effector in the gut brain axis [65,66]. While only produced at low levels in mono-cultures, it emerged as an abundant metabolite in a set of co-cultures, many of them involving *B. uniformis* (**SuppFig 6A**). Similarly, levels of indole-acetic acid, a key microbial metabolite affecting the immune system [25] and cancer therapy success [67], were boosted in several co-cultures involving *C. difficile* (**SuppFig 6B**). This observation is supported by recent findings that cross-feeding interactions are key to producing health-relevant indole metabolites [68,69].

Notable emergence was also detected for biogenic amines, including polyamines, which are produced from amino acid precursors, with important consequences for the host [62,63,70] (**Fig 4C**). Agmatine emerged in co-cultures of *E. coli* with *B. uniformis*, *S. copri*, *Parabacteroides merdae*, *A. rectalis* and *B. thetaiotaomicron*. Agmatine production is known to be induced by acid stress [71–73], consistent with media acidification by Bacteroidetes (**SuppFig 3B**). Here, putrescine emerged in co-cultures of *C. aerofaciens* with *E. coli*/*K. aerogenes* and the same conditions showed evidence for cross-feeding of ornithine produced by *C. aerofaciens* (**Fig 4D**). Ornithine is converted to putrescine by decarboxylation (**Fig 4E**) and it is therefore likely that the emergence of putrescine is due to cross-feeding of its precursor. Furthermore, histamine production by *K. aerogenes* was boosted above baseline by a set of partner species including *B. thetaiotaomicron* and *S. copri*. Cadaverine was produced by both *E. coli* and *K. aerogenes*, but this baseline production was not modulated in any co-culture. However, co-cultures with *S. copri* contained acetyl-cadaverine, suggesting that *S. copri* modifies cadaverine produced by *E. coli* and *K. aerogenes*. Acetylation neutralises the positive charge of polyamines with potential effects on bacterial physiology [74,75].

We next investigated if the changes in amine production can be explained by changes in enzyme abundance. We Identified no strong expression changes that correlate with the observed increases in amine production (**Fig 4F**), indicating that these phenomena originate from the post-translational or metabolic level. A notable exception is the ∼9-fold upregulation of *E. coli* SpeF in the presence of *C. aerofaciens* which correlated with an increase in putrescine in the co-culture. However, multiple other co-culture partners caused a similar (albeit weaker) upregulation of SpeF without concurrent putrescine production (**Fig 4E+F**). These results suggest that amine metabolism in the gut is emergent and highly dependent on the nutritional environment. While species such as *E. coli* and *K. aerogenes* clearly possess the genetic repertoire to produce a range of amines at high concentrations, this does not happen in all conditions, even if the precursors are abundant.

We also noted numerous interactions involving fermentation end products such as succinate, SCFA and BCFA (**SuppFig 6D**). Propionic acid was depleted in several co-cultures, but also surprisingly emerged in a small number of cases, three of which involving *L. saccharolytica* and a Bacteroidete. Succinate was depleted and butyrate emerged in several co-cultures of *Clostridia* with *Bacteroides*, in accordance with proteomic changes coherent with acetate and succinate cross-feeding (**Fig 3D**). Overall, these results underline the emergent and complex nature of communal metabolism and provide a framework for identifying potential metabolic interactions through comparative metabolomics.

### Narrow amino acid preferences indicate niche partitioning

Gut bacteria obtain amino acids from proteins/peptides in partially digested food [76] and from host-produced mucin [77] and in turn impact host amino acid status [78]. Many gut bacteria secrete proteases to digest peptides in the environment thereby releasing a mix of amino acids [79,80], yet gut bacteria were found to often have a narrow preference for amino acids [78]. Furthermore, amino acid auxotrophies are common among gut bacteria [81,82] and amino acids have been proposed as prebiotics [83]. Still, significant knowledge gaps remain around communal metabolism of amino acids. We therefore next focused on the metabolism of proteinogenic amino acids.

We determined the concentrations of free amino acids in the supernatant of our 608 co- and mono-culture samples. We found that mono-cultures of most species were depleted in none or very few amino acids (**Fig 5A**). While the uptake and intracellular metabolism of small peptides is not captured by these data, it is apparent that, in terms of free amino acids, gut bacteria have a narrow preference/requirement for amino acids. The closely related species *E. coli* and *K. aerogenes* had a similar preference profile and depleted lysine, serine, threonine and aspartate. However, the two closely related species *C. difficile* and *C. sporogenes* had distinct and broad profiles overlapping in serine and threonine only. Other species had notably narrow and distinct preference profiles, with e.g. *C. aerofaciens* selectively depleting arginine and *L. saccharolytica* depleting serine.

**Figure 5:**
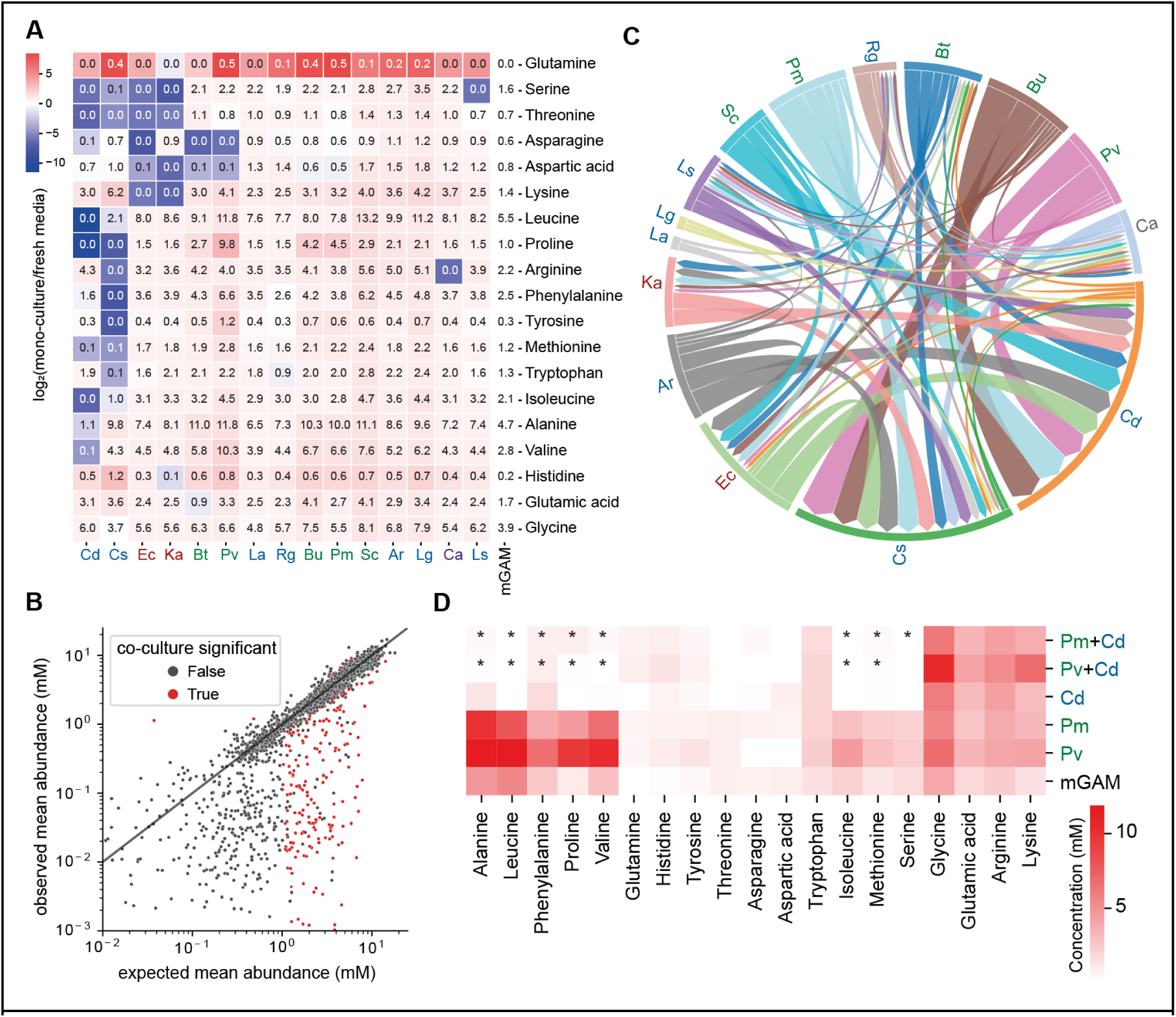
Narrow amino acid preferences result in niche partitioning and cooperative protein digestion. (A) Heatmap illustrating concentrations of free amino acids in mono-cultures versus fresh mGAM medium. Colours indicate the log_2_-transformed fold-change. Numbers in the heatmap indicate mean concentrations in mM (n≈16 biological replicates). (B) Comparison of expected amino acid concentrations based on mono-cultures, relative abundances and co-culture OD, with those observed in the co-cultures. Colours denote statistically significant interactions (abs(log_2_(fold-change)) >0.5, minimum predicted or observed concentration >1 mM). (C) Amino acid exchange network reconstructed using the data on amino acids unexpectedly depleted in co-cultures (identified in B) where donor and recipient species could be assigned (Methods). Ribbon widths reflect the number of exchanged amino acids. (D) Heatmap illustrating amino acid concentrations across mono- and co-cultures involving *C. difficile* and different Bacteroidetes (*S. copri, P. merdae* and *P. vulgatus*). Asterisks denote significant interactions (see panel B). The accumulation of a group of amino acids in the Bacteroidetes mono-culture compared to fresh mGAM medium but their absence in both *C. difficile* mono-culture and co-cultures indicates that these are likely consumed by *C. difficile*.

This pattern could indicate niche partitioning in the gut community, in a scenario where extracellular proteins/peptides are digested collectively, and specific released amino acids are used by different species. To investigate this, we compared observed amino acid concentrations in the co-cultures to expected values based on the absolute abundance of each species and the observed consumption/release pattern observed in mono-culture (**Fig 5B**). Out of 1976 potential amino acid-species pair interactions, we detected 187 cases where the observed concentrations differed from the expected (p_adj_<0.05, FDR-corrected two-sided Welch’s t-test, abs(log_2_(FC))>0.5, minimum predicted or observed concentration of 1 mM). In all but 5 cases, the amino acid decreased, suggesting that one of the species is consuming amino acids released by the other.

In 49 cases, we could predict the direction of amino acid flow based on the consumption/release patterns in the mono-culture, resulting in the metabolite exchange network shown in (**Fig 5C**). *K. aerogenes*, *E. coli*, *C. sporogenes*, *C. difficile* and *B. thetaiotaomicron* were overall receivers of amino acids. Leucine, lysine and phenylalanine acid stood out as being exchanged the most (12, 11 and 10 pairs respectively). Notably, the opportunistic pathogen *C. difficile* with a relatively broad amino acid uptake pattern received amino acids from a range of Bacteroidetes (**Fig 5D)**. Our data thus shows that amino acids obtained by proteolysis are divided amongst co-culture members with narrow-range requirements indicating both niche partitioning and nutrient sharing.

### Regulation of orthologous genes is highly divergent

Our analyses so far revealed complex interactions at the level of growth, protein expressions and metabolism. We next set out to leverage the data for functional and comparative genomics. Taking advantage of the evolutionary diversity of the dataset, we investigated the extent to which the regulatory responses of orthologues to partner species are conserved.

We used Orthofinder [84] to identify pairs of orthologous proteins and computed Pearson correlation between their abundance changes across co-culture conditions. In the cases of distantly related species pairs, we observed virtually no correlation between orthologue expression profiles (**Fig 6A**). Only closely related species showed notable correlation, e.g., *E. coli-K. aerogenes* (r=0.2), *L. acidophilus-L. gasseri* (r=0.17) and *Bacteroides* pairs. Exemplarily focusing on the species *P. vulgatus,* orthologue pairs with *B. uniformis* and *B. thetaiotaomicron* exhibited median correlations of 0.43 and 0.25 respectively (**Fig 6B**). The *B. uniformis-P. vulgatus* correlation was unexpectedly strong since the more closely related *B. uniformis-B. thetaiotaomicron* pair showed a weaker correlation. For the former pair, we noted distinct correlation patterns for different classes of proteins, with enzymes and transporters showing the tightest correlation (**Fig 6C**). For the other two Bacteroidetes, the correlation was lower (0.18) and for all other species it was below 0.1, indicating overall poor to no conservation of regulatory patterns.

**Figure 6:**
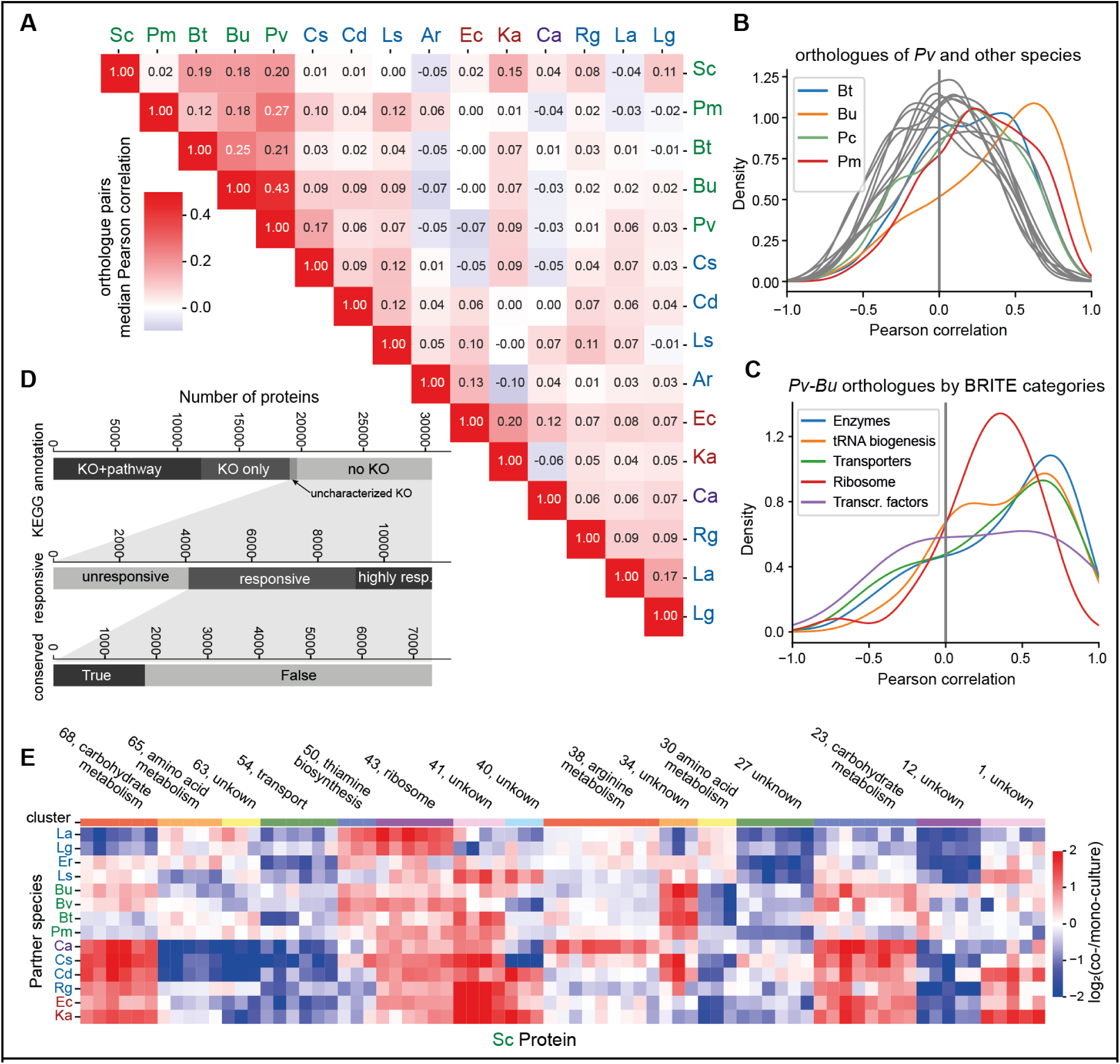
Evolutionary divergence of proteome responses and functional genomics. (A) Regulation of orthologs in response to other species is conserved only amongst closely related species. Median Pearson correlation coefficient between ortholog pairs of species pairs. The analysis was restricted to single-protein orthologue pairs, reliably detected proteins (<4 missing) and strongly changing proteins (sum(abs(log_2_(fc))) > 2). (B) Distributions of Pearson correlation coefficients of ortholog pairs between *P. vulgatus* and other species. Only closely related Bacteroidetes species (highlighted in colours) show a shift towards positively correlated expression profiles, indicating similar expression patterns. (C) Different functional categories of proteins displayed different levels of conservation in their expression profiles. Enzymes and transporters were the most similar, while ribosomal proteins were less similar and transcription factors displayed a broader distribution. (D) From top to bottom: Barplot illustrating status of KEGG annotation across dataset; responsiveness of proteins with uncharacterised KOs or without assigned KEGG KO; conservation for responsive and common responsive proteins without KEGG KO. Conservation was defined as eggNOG OG present in >70% of bacterial species. This highlights the very large number of responsive and common responsive proteins without any functional annotation. The approximately 1800 proteins therein with strong conservation across bacteria are priority unstudied genes in the context of communal gene expression. (E) Clustered heatmap showing a selection of highly regulated *S. copri* proteins. Several clusters are made up of proteins with similar functions indicating that co-expression clustering can inform about protein function. Many clusters contain only genes with no existing functional annotations. Hierarchical clustering was performed using Euclidean distance metric and the Ward method. The colour scale was clipped at -2,2. Numbers at the top indicate the cluster ID (see <u>Supplementary Data 8</u> for all clustering results).

While some studies have compared expression profiles of orthologous genes at the transcriptome level, e.g. [85–87], the evolution of protein abundances remains understudied. Our results indicate that the condition-specific regulation of protein expression is highly variable and even closely related species execute different expression profiles in response to the same partner species. This could point to both functional diversification at the protein level (homologous proteins performing different functions and therefore being regulated differently) and at the species level (species reacting differently to a similar environment as it affects them differently due to metabolic or cellular differences). The divergent regulation of orthologous proteins in gut bacteria thus highlights the need to account for inter-species interactions in understanding protein and organismal functions.

### Poorly characterised proteins are responsive in co-culture and cluster according to function

The interpretation of gene expression data depends on the availability and quality of functional annotation of proteins. However, a large fraction of bacterial genes, even in relatively well studied gut bacterial species, remains functionally unannotated. In our set of species, 50% of proteins in the genome and 37% of the measured proteome were not annotated to a KEGG Orthology (KO) term or were annotated to an uncharacterised KO (**Fig 6D**). This ‘genetic dark matter’ presents an enormous blind spot in omics studies [1,88,89]. 64% of these poorly characterised proteins were responsive (at least one hit) or common responsive (at least 4 hits) in our dataset. Out of the poorly annotated but responsive proteins, 24% (n=1781) were conserved across bacteria (data from eggNOG, Methods), indicating a key physiological role in the community context. This approach therefore enables the prioritisation of unstudied genes for further, focused experiments.

We next used an unsupervised clustering approach to group proteins with similar expression profiles. Out of 1035 clusters across all species, 185 were enriched in at least one KEGG pathway, and 713 clusters contained at least one gene pair that shared a genomic neighbourhood. These findings indicate that this approach can identify groups of proteins with related biological function.

To explore the use of systematic co-culture proteomics for functional genomics, we focused on *S. copri*, an abundant yet understudied gut microbe negatively correlated with western diet [90]. From 75 clusters in total, we selected a subset of smaller clusters with highly coherent and diverse expression profiles (**Fig 6E, SuppFig 7**). Some of these contained genes clearly attributable to specific functions, such as cluster 38 which contained arginine biosynthesis genes upregulated in co-culture with *C. aerofaciens*, cluster 50 which contained thiamine biosynthesis genes upregulated in a range of co-cultures including with *C. sporogenes*, or cluster 43 which contained ribosomal proteins. In some clusters, annotated proteins clustered with uncharacterised ones suggesting shared function. E.g., cluster 68 contained several poorly characterised carbohydrate metabolism genes together with xylan-metabolising enzymes in the same genomic neighbourhood. However, many clusters showed no existing high-quality functional annotation for any proteins. E.g., cluster 41 contains 4 neighbouring genes and was strongly upregulated in co-culture with *E. coli*, *K. aerogenes*, *C. sporogenes* and *R. gnavus*. This case study illustrates the potential of using systematic (meta-)proteomics for co-expression analysis and functional genomics. Full information on clusters across all species is available in <u>Supplementary Data 8</u>. An interactive shiny app to explore the clustered expression profiles of all species is available at: https://stephan-kamrad.shinyapps.io/co-culture_proteomes/.

## Discussion

A key aspect of community ecology is the emergence of collective functions. Our systematic study enabled expansion of the cases of emergent metabolism previously observed in gut bacterial communities, e.g. in the case of bile acids [91] or carbohydrate metabolism [18]. Our findings include sequential metabolism, e.g., acetyl-cadaverine production in *E. coli*-*S. copri* co-cultures, and activation of a secondary metabolic pathway, e.g., the stimulation of agmatine in *E. coli* by some Bacteroidetes. We also identify candidate cross-fed metabolites as those depleted in co-cultures of species with complementary production-consumption profiles, e.g., ornithine being specifically depleted in *C. aerofaciens*-*K. aerogenes* co-cultures. Together, we demonstrate that metabolomics of combinatorial co-cultures is a promising strategy for identifying metabolic interactions and co-metabolism.

The observed metabolic interactions are closely intertwined with the proteome, evident in the strong enrichment of enzymes and transporters among the responsive proteins. Some metabolic changes correlated directly with enzyme abundances, e.g., arginine biosynthesis being upregulated in the presence of the arginine consumer *C. aerofaciens*. In other cases, metabolic changes appear to occur without changes in enzyme abundance, e.g., polyamine synthesis. This is congruent with pathways being regulated by either enzyme availability (enzyme control) or by metabolite availability (metabolic control). Thus, proteome and metabolome serve as complementary readouts, with enzyme regulation data indicative of metabolic changes directed by the cellular regulatory machinery. In the case of *C. difficile* interactions with *Bacteroides*, our comprehensive dataset captures interactions relating to starch metabolism, SCFA and succinate cross-feeding, sharing of amino acids and modulation of pH. This multiplicity of interactions, each depending on nutrient availability/media composition, explains the diversity of interaction types observed in these species in the literature [14,55–58,92]. Our results thus exemplify how the functional manifestation of the genetic repertoire is plastic and dependent on the community context.

Omics of *in vitro* cultures have resulted in deep insights into the biology of a few model bacteria such as *E. coli* and *Bacillus subtilis*. However, the vast majority of abundant and prevalent species remains massively understudied. Beyond relatively well annotated protein classes like enzymes, the lack of functional annotation represents a major hurdle to the interpretation of omics data from non-model bacterial species and communities including them. In our dataset, more than a third of genes were not annotated to a KEGG KO term. If large proteomic datasets are available, co-variation analysis can be a powerful way of gene functional annotation, which at least in human [93] and yeast [94], outperforms other strategies of functional annotation of unknown protein function, and is a particularly attractive strategy for annotating gene function in species that are difficult to manipulate genetically. We have here shown that systematic co-culture proteomics cluster into functionally enriched sets of proteins. At the same time, our case study of *S. copri* shows that many strongly and specifically regulated protein clusters are entirely un-annotated to date, thereby identifying priority unstudied genes in the community context. Our co-culture proteomics data is thus a useful complementation to other current systematic functional genomics efforts in the microbiome space, e.g. involving forward genetic screens through transposon mutagenesis [95,96].

Taken together, our data opens a molecular window into bacterial interactions with implications for functional genomics and for engineering community metabolism.

## Materials and Methods

### Cultivation of bacteria and sample collection

Bacterial isolates were obtained from culture collections (<u>Supplementary Data 1</u>). All cultures were grown statically at 37°C in an anaerobic polyvinyl chamber (Coy Instruments) filled with 2.5% H_2_, 12% CO_2_ and balance N_2_. Cultures were grown in modified Gifu anaerobic broth (mGAM), produced by Nissui Pharmaceuticals and obtained from HyServe, prepared according to the instructions from the manufacturer and sterilised by autoclaving. From glycerol-preserved cryostocks, 10 mL cultures were grown in screw-top tubes for one or two days. Cultures were then diluted 100-fold into 10 mL of fresh media and incubated again for one day. The optical density at 600 nm (OD) of each culture was measured and a dilution was prepared. Using a 96-channel pipetting robot, diluted cultures were combined in all possible pairs in 96-well deep well plates at an initial OD of 0.05 and a volume of 1.4 mL. Columns 9-12 of each plate were used for mono-culture controls which had the same volume and initial OD. Plates were sealed with a breathable film and incubated for 24 h.

Plates were then removed from the anaerobic chamber, cultures were mixed well and 100 μL were transferred into a fresh 96-well plate which was used to measure the OD with a microplate reader. The remaining cultures in the deep-well plates were then centrifuged (5 mins, 3,200 g) and three times 80 μL of supernatant was collected for metabolomic analysis. The rest of the supernatant was discarded and 1 mL of water was added to wash off residual media from the cell pellet. Plates were shaken briefly before centrifuging again with the same settings. The supernatant was discarded and cell pellets frozen at -80°C until processing for proteomics.

### Proteomics sample preparation

Proteomics samples were prepared based on a previously established high-throughput protocol [94], with a few key modifications. An aqueous lysis buffer with the following composition freshly prepared: 7 M urea, 100 mM ammonium bicarbonate, 24 mM Tris-HCl, 2.4 mM MgCl_2_, 122 units/mL benzonase (Sigma-Adrich). A small amount of glass beads and 200 μL of lysis buffer were added to each pellet and the plates were sealed with rubber mats. Cells were lysed using beat beating (2x 5 mins, 1500 rpm), followed by a brief centrifugation and incubation for 20 min at 37°C in a water bath (for benzonase digestion). For reduction-alkylation, 20 μL of 55 mM dithiothreitol were added, plates were mixed briefly and incubated at 30°C for 1h, followed by 20 μL of 120 mM iodoacetamide, brief mixing and 30 min at room temperature in the dark. 450 μL of 0.1 M ammonium bicarbonate was added, plates were mixed briefly and centrifuged for 5 min at 3,220 g to clear the extract from cell debris.

Using an automated workflow on a liquid handling robot and the previously obtained OD measurements of each culture, 0.72 OD units of extract were transferred to a fresh deep well plate and adjusted to a final volume of 500 μL using an aqueous solution of 2 M urea and 0.1 M ammonium bicarbonate. This ensured an approximately even input biomass for all samples, while minimising the time that live bacterial cells are handled aerobically before lysis and denaturation.

For digestion, 10 μL of trypsin/LysC mix (Promega, Cat# V5072, prepared according to instructions from the manufacturer) were added to each sample with subsequent incubation at 37°C with shaking for 17 h. Digestion was stopped by adding 25 μL of 25% formic acid. Solid phase extraction in 96-well format was used to clean up the samples. SPE plates (BioPureSPE PROTO C18 MACRO, Nest Group, Cat# HNS S18V-L) were first conditioned with methanol, followed by twice buffer A (50% acetonitrile in water) and twice buffer B (water with 0.1% formic acid), using a volume of 200 μL per well throughout. The entire digest was then loaded and washed with three times buffer B before eluting with three times 110 μL of buffer A. Samples were dried in a concentrator (Savant SPD300DDA) at 35°C for approximately 6 h. Samples were reconstituted in 40 μL of buffer B and peptide concentrations were determined using a fluorometric assay (Pierce Quantitative Fluorometric Peptide Assay, ThermoFisher, used according to manufacturer’s instructions).

### Proteomics data acquisition

LC-MS analysis was performed on an Evosep One system coupled to a Bruker timsTOF Pro 1 mass spectrometer. 200 ng of peptides were loaded on Evotip Pure tips according to the manufacturer’s protocol. Liquid chromatography was carried out using the Evosep 100 SPD LC method (11.5 minutes gradient) with an EV1109 performance column (ReproSil Saphir C18, 8 cm x 150 µm, 1.5 µm beads by Dr Maisch) at 40°C, coupled to a 10 µm ZDV (ZeroDeadVolume) captive Spray Emitter. Data was collected over an m/z range of 100 to 1700 for MS on the timsTOF Pro instrument using an accumulation and ramp time of 100 ms. The ion mobility range was set 0.85 to 1.27 Vs cm^-2^. DIA-PASEF scans used a cycle time of 0.95 s. Mobility windows included 21 mass steps per cycle with a 25 Da mass width. The collision energy was decreased as a function of the ion mobility from 20 eV (1/K_0_ = 60 Vs cm^-2^) to 59 eV (1/K_0_= 1.6 Vs cm^-2^).

### Proteomics data analysis

Proteomics raw were analysed in two stages using DIA-NN v1.8.2 beta 27 [45,97]. In stage 1, all samples were searched against an *in silico* predicted library generated from the proteome fasta files of all species (obtained from Uniprot [98]). DIA-NN was called from the command line using the following options: --qvalue 0.01 --matrices --gen-spec-lib --predictor --fasta-search --min-fr-mz 200 -- max-fr-mz 1800 --met-excision --cut K*,R* --missed-cleavages 1 --min-pep-len 7 --max-pep-len 30 -- min-pr-mz 300 --max-pr-mz 1800 --min-pr-charge 2 --max-pr-charge 3 --unimod4 --window 18 --mass-acc 15 --mass-acc-ms1 15 --smart-profiling --pg-level 1

Data from stage 1 was used to assess sample quality. Out of 768 samples for which data was acquired, 106 samples were excluded because they had fewer than 4000 precursor identifications or more than 10% of identified precursors were originating from species other than those expected in the sample. We furthermore excluded all samples containing the species *Bifidobacterium longum subspecies longum* as very few precursors from this species were detected across the dataset, most likely due to the low growth rate of this species and a contamination of the pre-culture. Another 8 samples were excluded as these showed abnormally high ODs compared to the other replicates, likely indicating contamination. A total of 608 samples were included for further analysis.

In stage 2, one DIA-NN call was performed per species, searching all samples containing the species against a predicted spectral library for that species only. I.e. each co-culture sample was analysed in two separate DIA-NN calls, one for each species. Options were as above, except that we enabled ‘Match Between Runs’, which performs a two-pass search, the second of which with a subsetted library of precursors detected in at least one sample. Peptide reports in parquet format were then further processed in Python and R by removing precursors with Q.Value>0.01, Global.Q.Value>0.01 and Quantity.Quality<0.75. Non-proteotypic peptides were removed (including those matching to more than one protein within the species, as well as those matching to a protein from another species in the sample). We also removed a small number of peptides which were detected consistently (in more than 3 cases) in the mono-cultures of the other species in the sample. Finally, precursors were excluded if their coefficient of variation across the mono-culture replicates was greater than 0.5, indicating strong technical or biological noise.

After precursor filtering, the normalised quantUMS [99] precursor quantities computed by DIA-NN (column ‘Precursor.Normalised’) were log_2_-transformed and re-normalised using cyclic loess normalisation as implemented in limma [100] using default options. Protein quantities were computed with the maxLFQ algorithm [101] as implemented in die DIA-NN R package after transforming back to normal (i.e. not log_2_) scale. A small number of proteins with a CV>0.5 across the mono-culture replicates were excluded. Protein-level summary statistics were computed and statistical analysis was performed with limma by fitting a linear model to the data (using the lmFit function), defining contrasts as each co-culture versus the mono-culture, and using empirical Bayes to obtain moderated t-statistics where standard errors were moderated towards a trend using eBayes(trend=True). For proteins which were detected in a co-culture but not any of the mono-cultures, a fold change of 2 and an adjusted p-value of 0.049 was assumed.

### Determination of species abundances

We used the peptide report from stage 1 (above), filtered for proteotypic peptides (across all species) and q-value and quantity.quality filters as before. For each species, highly complete precursors were selected (identified in >90% of samples) and missing values imputed using median imputation. The sum of non-normalised precursor quantities (column Precursor.Quantity) was used as proxy of relative species abundance which was divided by the same value obtained for the mono-culture of that species to correct for the fact that an identical amount of injected peptides will produce different intensity sums for each species (e.g. due to the quality of the annotated proteome or physicochemical peptide properties). For each co-culture sample, the quantities obtained for each species were divided by the sum of both to obtain relative abundances for each sample. These were then scaled by the total OD600 of the co-culture to obtain absolute abundances of each species in the co-culture in OD units.

### pH measurements

High throughput colorimetric pH estimations were conducted as described [102], with slight modifications. In brief, a 21-point pH calibration curve between pH 4.03 and 8.08 was prepared in mGAM and the pH of each calibrant was measured using a micro pH meter (ThermoScientific 8220BNWP). 10 μL of 10 mg/mL litmus dye was added to 50 μL of each calibrant as well as supernatant samples in half-area flat-bottom plates (Greiner, Cat#675801). Full absorption spectra (350 nm to 800 nm, with 5 nm step-size) of three replicates of each point in the calibration curve were acquired on a Varioskan (ThermoScientific VL0000D0) and used to determine the isosbestic point (wavelength corresponding to minimum absorption variance across calibrants) and the wavelength corresponding to maximum absorbance variance across the calibrants). Absorbance measurements at these two wavelengths (430 nm and 585 nm, respectively) were acquired for all calibrants and test samples. A sigmoidal curve fitted using the absorbance ratio (abs. at 430 nm/ abs. at 585 nm) and calibrant pH was used to estimate supernatant sample pH.

### Prediction of responsive proteome fraction

The multiple linear regression model was built with scikit-learn. Features were centred and scaled using scale from sklearn.preprocessing; Linear regression was performed with LinearRegression from sklearn.linear_model; cross-validation was performed using LeaveOneOut from sklearn.model_selection. Default settings were used throughout.

### Metadata and genomic analysis

#### Functional annotation

KEGG annotations for protein sequences were retrieved with the GHOST Koala tools via the web interface. EggNOG annotations for protein sequences were obtained using the eggNOG mapper v 2.1.12 [103] via the web interface. Genomes were retrieved from sources listed in <u>Supplementary Data 1</u> and gene locations were identified using tblastn [104,105]. The conservation of proteins across bacteria was computed using data downloaded from eggNOG (v 5.0) [103]. Specifically, the table of proteins assigned to each eggNOG OG (/download/eggnog_5.0/per_tax_level/2/2_members.tsv.gz) was used to count the number of species in which a given OG is present and this number was divided by the total number of bacteria in the database (5090, see file e5.taxid_info.tsv in the same folder). A protein was considered conserved if it was present in more than 70% of species. Metabolic models of the organisms of interest were drafted with CarveMe v.1.6.1 [106] using the proteome as input.

#### Enrichment analyses

Fisher’s exact test as implemented in the scipy.stats module was used to test for significant enrichment. All measured proteins (of the respective species) were used as the background set. Enrichment was quantified as (hit proteins annotated to term/all hit proteins) / (background proteins annotated to term/background proteins).

#### Clustering

Clustering was performed on log_2_-transformed fold-change data (co-/mono-culture). For clustering, proteins with more than 3 missing values were removed. We also excluded proteins which showed very little response across conditions (sum(abs(log_2_(fold-change))) < 2). Remaining missing values were imputed with 0. Hierarchical clustering was performed with Euclidean distance metric and Ward linkage method, as implemented in AgglomerativeClustering from the scikit-learn sklearn.cluster module. The number of clusters was set to the number of proteins divided by 20.

#### Orthologue analysis

Orthologues were identified using OrthoFinder v 2.5.5 [84] with default settings using protein sequences as input. To compute distributions of orthologue expression profile correlations, the OrthoFinder output for each species pair was filtered to only include single protein ortho-groups and only those where proteomics data was available for both proteins. Pearson correlation was used to compute the similarity of two orthologues’ expression profiles across co-culture conditions. Phylogenetic distances between species were computed based on the previously generated tree using the cophenetic.phylo function of the ape R package [107].

### Quantitative flow cytometry

Cells were stained to distinguish gram-positive and -negative cells by flow cytometry as described previously [108]. Cells from 1 mL of overnight culture were collected by centrifugation (5 min, 8,000 g), washed in 1 mL 1 M KCl, resuspended in 400 µL of 4% formaldehyde diluted in 1 M KCl, and incubated on ice for 30 minutes to fix the cells. The cell suspension was then centrifuged and the pellet was washed once with 1 M KCl. Cells were resuspended in 300 µL of 1 M KCl and stored at 4 °C until analysis. For staining, the samples were first diluted to OD 0.5 in 300 µL of 1 M KCl, followed by addition of 4 µg/mL Vancomycin BODIPY FL Conjugate (Invitrogen, Cat# V34850) and 15 minutes incubation at 30 °C in the dark. Cells were collected by centrifugation and resuspended in 300 µL of PBS in polystyrene tubes for flow cytometry.

Samples were run on a BD LSRFortessa cell analyser using the BD FACSDiva software and with the following settings: slow speed acquisition mode, laser 488 nm and the Blue650, FSC and SSC filter for acquisition. Fcs files were analysed with FlowJo (v 10.9.0) gating the cell population in the FSC-H/SSC-H plot, following a singlet gating on SSC-H/SSC-A plot and gating in the in a Blue530-H/FSC-H dimensions for discriminating gram positive (*C. difficile*) and gram negative (Bacteroidetes, *E. coli*) bacterial cells (**SuppFig 2**). The same samples were used for cell counting using the bacteria counting kit (Invitrogen, Cat# B7277) following the manufacturer’s protocol. Fixed cells were diluted in order to obtain a sufficient signal in the bead gate upon flow cytometry analysis and stained with Sytox so that the cells could be discriminated from the beads using the Blue530/30 and FSC filters (**SuppFig 2**).

### *B. thetaiotaomicron* sus mutants

*Escherichia coli* S17λpir was grown in Miller’s lysogeny broth (LB) (Corning) supplemented with ampicillin (100 μg/mL) when required and incubated at 37°C with 180 rpm shaking. Genetic constructions were made in *B. thetaiotaomicron* VPI-5482*Δtdk* background, developed for a two-step selection procedure of unmarked gene deletion by allelic exchange, as previously described [109]. All primers used in this study are listed in **SuppFig 5A**. Mutants were generated via allelic exchange using the suicide vector pLGB13 [110]. Approximately 500 bp regions flanking the target gene were amplified by PCR using Phusion Flash High-Fidelity PCR Master Mix (Thermo Fisher Scientific), and assembled with the plasmid backbone using Gibson assembly. The reaction mix, containing ISO buffer, T5 exonuclease, Phusion HF polymerase, and Taq DNA ligase, was incubated at 50°C for 35 minutes. The assembled plasmid was then introduced into *E. coli* S17 λpir, which served as the conjugation donor. For conjugation, exponentially growing cultures of donor and recipient were mixed at a 2:1 ratio and spotted on BHIS agar (Brain-Heart Infusion medium supplemented with 5 mg/L hemin, 2 g/L NaCOH_3_, and 1 g/L cysteine), followed by overnight incubation at 37°C under aerobic conditions. The following day, the mix was plated on BHIS agar supplemented with erythromycin (15 µg/mL) to select for *B. thetaiotaomicron* transconjugants that had undergone the first recombination, and gentamicin (200 µg/mL) to inhibit donor *E. coli* growth. Resulting colonies were cultured overnight in antibiotic-free BHIS medium to facilitate plasmid loss, then plated on BHIS agar containing anhydrotetracycline to counterselect against cells retaining the vector. Candidate deletion mutants were verified by colony PCR using flanking primers, followed by Sanger sequencing. Growth on single carbon sources was tested with *B. theta* WT and genetic constructs deficient in hydrolase enzyme (*B. theta* ΔSusG) and starch-binding protein (*B. theta* ΔSusD), in M9 defined media containing single carbon source, or glucose or amylopectin, a branched polymer of glucose (**SuppFig 5B**). M9 defined medium consisted of M9 salts (6 g/L Na_2_HPO_4_, 3 g/L KH_2_PO_4_, 0.5 g/L NaCl, 1 g/L NH4Cl), 0.246 g/L MgSO_4_.7H2O, 0.014 g/L CaCl_2_.2H2O, 50 mg/L cysteine, 5 mg/L hemin, 2.5 µg/L vitamin K3, 5 µg/L vitamin B12, 2 mg/L FeSO_4_ 7H_2_O, and 1 g/L carbon source (glucose or amylopectin). Amylopectin from maize was autoclaved and dialysed using 3.5 kDa MW membranes prior to use (Slide-A-Lyzer Dialysis Cassettes, ThermoScientific).

### Metabolomics sample preparation

Supernatant was collected from cultures in a fresh 96-well plate and stored at -80°C until further processing. For biogenic analysis, supernatants were diluted 1:10 in water in a final volume of 80 µL; for amino acid analysis, supernatants were diluted 1:100 in water in a final volume of 80 µL; for all other LC-MS methods, 80 µL of undiluted supernatant was used. (Diluted) supernatants were extracted by protein crash by adding 120 µL of extraction buffer (1:1 acetonitrile:methanol with 0.1% formic and 20 µM of each amoxicillin, caffeine, ibuprofen and donepezil as internal standards), mixing briefly by shaking (15 s, 1500 rpm), incubating at 4°C for approximately 30 min and centrifuging (5 mins, 3,200 g, 4°C). 15 µL of cleared extract was removed into a fresh 384-well PCR plate for LC-MS analysis.

### Metabolomics data acquisition

All metabolomics measurements were performed on an Agilent Infinity 1290 LC coupled to an Agilent 6470B triple quadrupole mass spectrometer with JetStream ion source. Multiple reaction monitoring (MRM) was used to monitor compound-specific precursor-fragment transitions (usually two per compound). A dilution series of a mixed analytical standard, water blanks, media blanks and a QC sample (obtained from an *E. coli* culture, spiked with standard solution if required) were injected in regular intervals. Samples were injected in random order. Amino acids were measured as described previously [111,112], using a low-pH hydrophilic interaction chromatography (HILIC) method. Short- and branched-chain fatty acids were measured without derivatisation on a porous graphitic carbon column (Hypercarb, ThermoFisher) using the method described in ref [113]. Organic acids were measured using a high-pH HILIC method [114]. Amines were measured using a shortened low-pH HILIC method adapted from ref [115]. Tryptophan metabolites and various other non-polar gut-specific metabolites were measured using a custom reverse phase method. Further details, including a list of all transitions and method parameters is provided in <u>Supplementary Data 5</u>.

### Metabolomics data analysis

Raw data were analysed using MassHunter Quantitative Analysis. Compounds which were not detected in any of the samples or which had bad calibration curves, unacceptably high background signal or bad peak shapes were excluded from further analysis. Concentrations were estimated from peak areas using calibration curves obtained from serially diluted mixed standards. The data was further processed using custom scripts. Only samples matching those used in the proteomics experiment were taken forward for analysis, i.e. samples which were removed from the proteomics analysis due to suspected contamination or other reasons were also removed here.

### Metabolomics statistical analysis

Identification of compounds produced by mono-cultures - To identify which species produces/consumes metabolites in mono-culture, significant differences in the mean concentrations across biological replicates between mono-culture and fresh mGAM were identified using FDR-corrected two-sided Welch’s t-test. For proteinogenic amino acids significance was defined as p_adj_<0.01 and abs(log_2_(fold-change)) >1; for other compounds it was defined as p_adj_<0.01, fold-change >5 and minimum concentration in mono-culture > 500 μM for SCFA and >50 μM for all other compounds.

Interaction analysis - For each sample, metabolite concentrations were subtracted by the mean value obtained for fresh media to obtain concentration changes. For each co-culture sample the expected metabolite concentration was determined by randomly sampling one of the mono-culture concentrations, multiplying by the respective relative species abundance (absolute abundance in co- /mono-culture) and adding these to the baseline concentration in fresh media. Independent, two-sided Student’s t-test with FDR correction was used to compare expected to observed concentrations across the ∼4 replicates. A significant interaction was called if p_adj_<0.05, abs(log_2_(fold-change))>0.5 and minimum observed or predicted concentration > 50 µM (1 mM for amino acids). Amino acid donors and recipients (**Fig 6C**) were assigned based on analysis described in the paragraph above.

## Data Availability

The following supplementary data is provided with this manuscript:

Supplementary Data 1 - Bacterial strains

Supplementary Data 2 - Proteomics summary statistics

Supplementary Data 3 - Ecological interaction types

Supplementary Data 4 - Predicting the fraction of responsive proteins

Supplementary Data 5 - Metabolomics method details

Supplementary Data 6 - Metabolomics summary statistics

Supplementary Data 7 - Emergent and cross-fed metabolites

Supplementary Data 8 - Gene clusters and functional annotation

The mass spectrometry proteomics data have been deposited to the ProteomeXchange Consortium (http://proteomecentral.proteomexchange.org) via the PRIDE partner repository [116] with the dataset identifier PXD055395.

The proteomics data can be explored in our interactive web app: https://stephan-kamrad.shinyapps.io/co-culture_proteomes/

Targeted metabolomics raw data has been deposited to Mendeley Data with DOI 10.17632/8wsm6tkh6n.1, accessible at https://data.mendeley.com/datasets/8wsm6tkh6n/1

## Competing Interest Statement

The authors declare no competing interests.

## Author Contributions

SK, KP and MR conceptualised the study. SK, SB, IC, DS, RB, SKA and SM performed the experiments. SK analysed the data and wrote the original manuscript draft. AB, NIvdB, and RG contributed to data analysis. KP, MR and MM supervised the study. All authors reviewed the manuscript.

## Acknowledgements

The authors would like to thank Vadim Demichev, Vadim Fartzdinov, Oliver Lemke, Daniela Ludwig, Lukasz Szwyriel, Lisa Kahl, Rahma Ute and Agatha Niewienda (Institute of Biochemistry, Charite University Medicine, Berlin, Germany) for help and advice with data acquisition and analysis. We would like to thank Samuel Shum (MRC Toxicology Unit, Cambridge, UK) for help with data management. We would like to thank Lucia Pinon Giraldez and Liam Cocker for advice and assistance with flow cytometry analysis. Nonantzin Beristain-Covarrubias (MRC Toxicology Unit, Cambridge, UK) provided materials. We thank the Genetics of Biofilms Unit (Institut Pasteur, Paris) for their guidance in designing the Bacteroides deletion mutants. This project has received funding from the European Research Council (ERC) under the European Union’s Horizon 2020 research and innovation programme (grant no. 866028) and from the UK Medical Research Council (project no. MC_UU_00025/11). SM was supported by HFSP award LT0018/2023. DS acknowledges the Swiss National Science Foundation Grant (P500PB_211100).

## Supplementary Figures

**Supplementary Figure 1:**
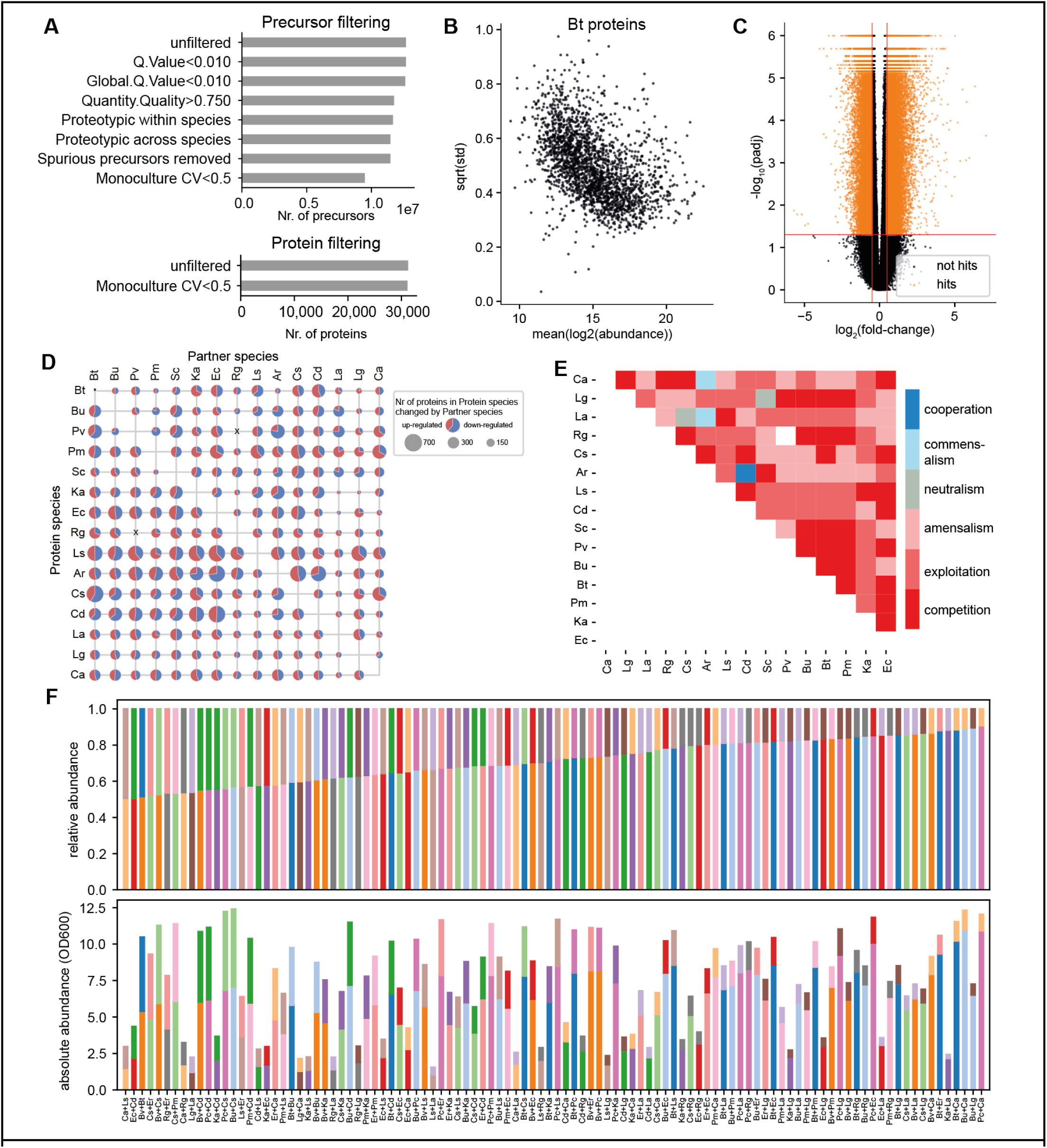
Proteomics data processing and ecological interactions. (A) Illustration of precursor- and protein-level filtering. Bar widths reflect the number of remaining precursors/proteins after each filtering step. This plot aggregates data from the 15 individual DIA-NN runs (one per species). Each run was processed separately. Precursors were filtered as shown from top to bottom (also see Methods). The filtered precursors were then re-normalised across samples and protein quantities were computed with maxLFQ. Proteins were filtered based on mono-culture CV only. (B) Differential abundance was assessed using empirical Bayes statistics as implemented in limma. This method squeezes the variance towards a global abundance versus variance trend, which is illustrated here exemplarily for the species *B. thetaiotaomicron*. (C) A fold-change cut-off of abs(log_2_(co-/mono-culture) and a significance cut-off of p_adj_<0.05 (FDR corrected, species-wise) was applied to the differential abundance statistic as illustrated in this volcano plot which aggregates data for all 15 species. 14.07% of the tested interactions were significant. (D) Number of hits in each co-culture. (E) Ecological interaction types for all co-cultures. (F) Relative abundance (top) and absolute abundances (relative abundances multiplied by co-culture OD, bottom) for all 104 co-culture pairs.

**Supplementary Figure 2.**
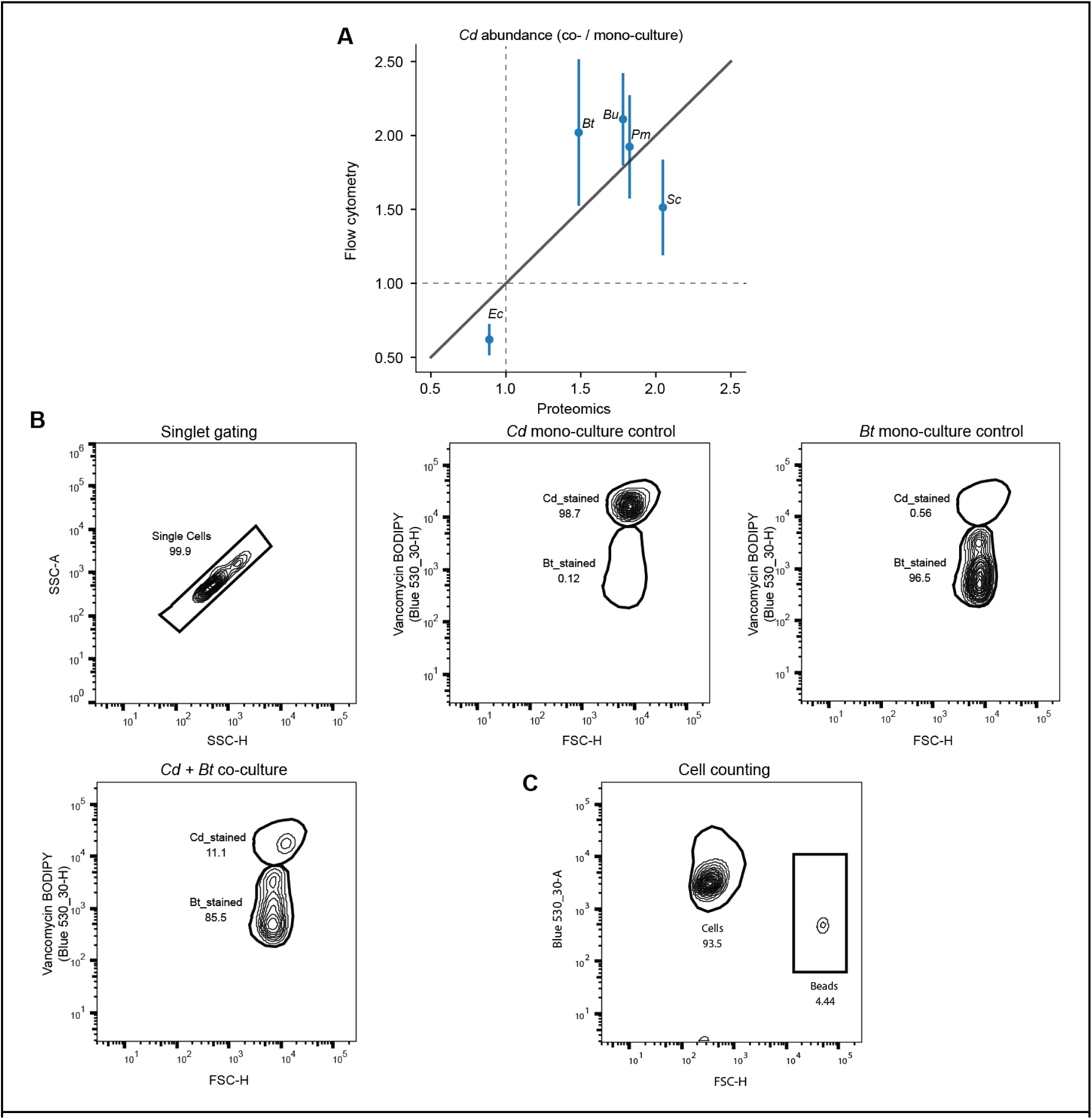
Quantitative flow cytometry. (A) Ratio of abundance of *C. difficile* in co- versus mono-culture with 5 selected partner species obtained via main proteomics experiment (x-axis, subset of data shown in **Fig 1J**) and quantitative flow cytometry (y-axis). Flow cytometry was performed on n=3 biological replicates, errorbars indicate the standard deviation. This independent measurement confirms that the absolute abundance of *C. difficile* increases in co-culture with the selected Bacteroidetes but decreases in co-culture with *E. coli*, as indicated in the main proteomics experiment. (B) Representative plots illustrating the gating strategy used for relative quantification of species via stain which distinguishes gram positive/negative cells (vancomycin-biodipy conjugate). (C) Representative plots illustrating the gating strategy used for absolute quantification of bacterial cells via bead spike in.

**Supplementary Figure 3.**
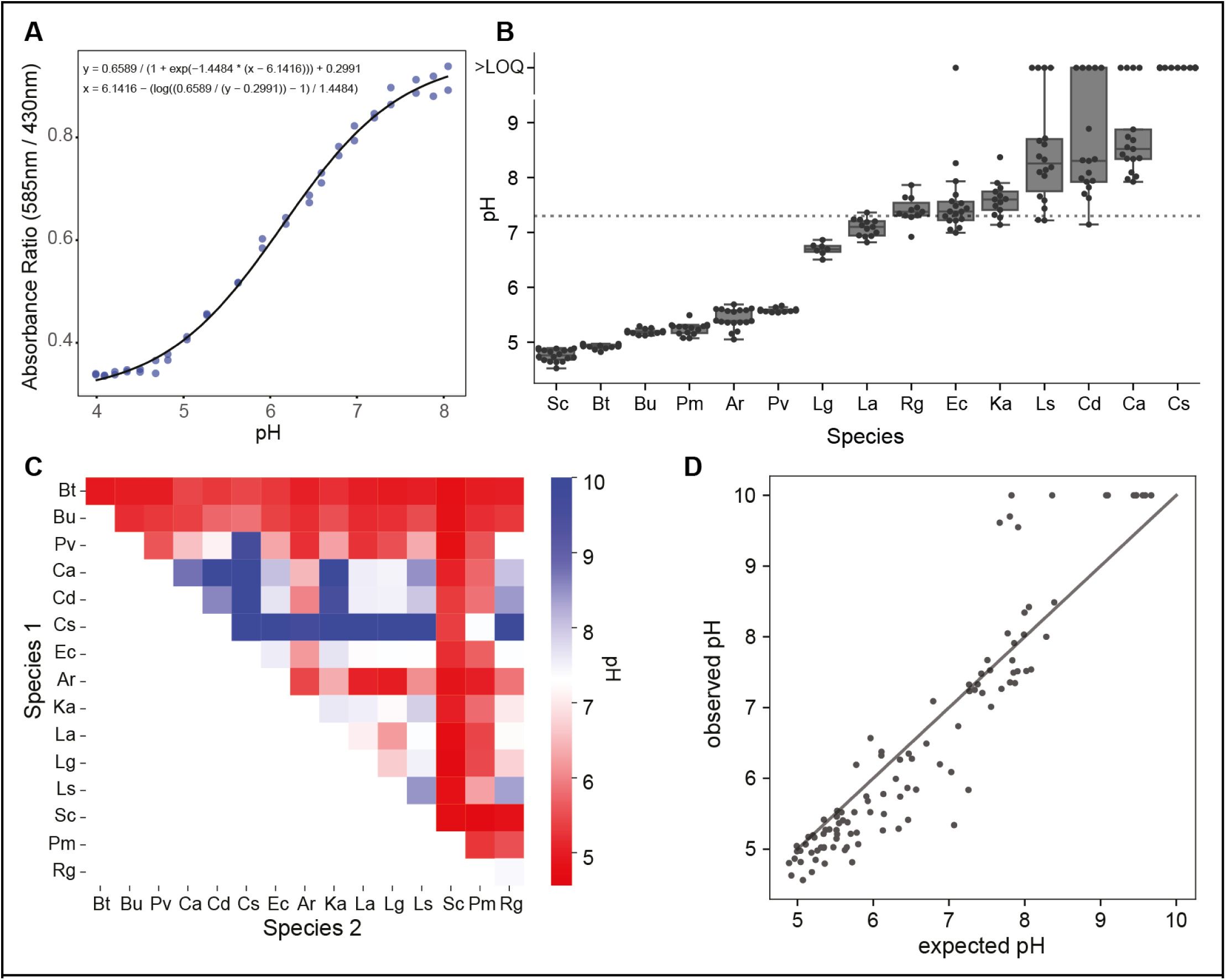
pH. (A) Calibration curve of high-throughput pH assay using litmus dye, (B) Mono-culture pH values. LOQ = limit of quantification. The dotted line indicates the pH of fresh media. (C) Co-culture pH values. (D) Observed versus expected (average of mono-culture pH, weighted by relative abundance of species in co-culture).

**Supplementary Figure 4.**
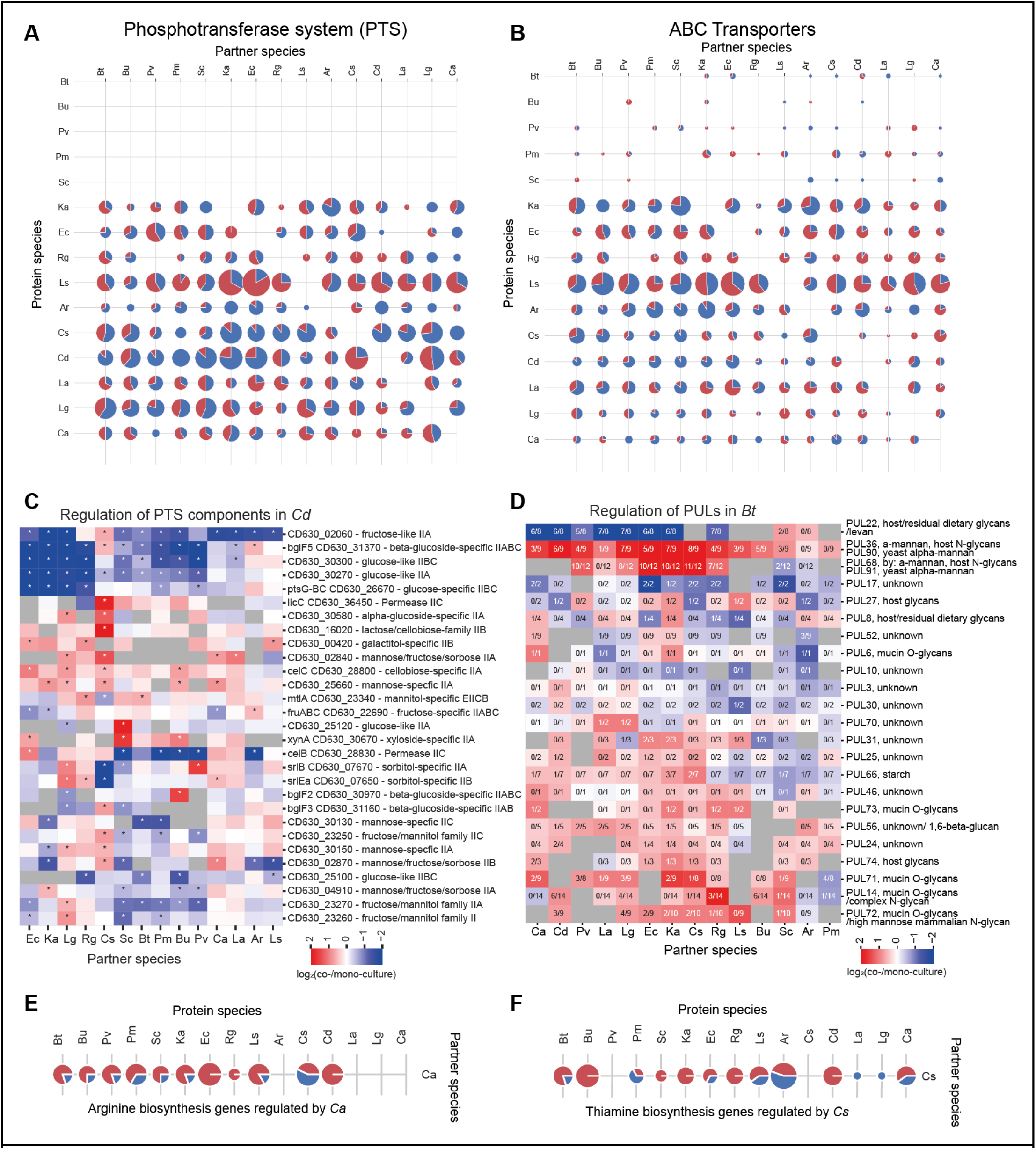
Differential protein abundance by functional groups. (A) Number of up- (red) and down-regulated (blue) proteins annotated to the phosphotransferase system (PTS) in each protein species in the presence of each partner species. The PTS system is absent in *Bacteroides*. (B) Number of up- (red) and down-regulated (blue) proteins annotated to the ATP-binding cassette transporters (ABC) in each protein species in the presence of each partner species. *L. saccharolytica* has an extraordinarily high number of (regulated) ABC transporters. (C) Regulation of PTS system components in *C. difficile*. Asterisks indicate significant interactions. (D) Regulation of polysaccharide utilising loci (PULs) in *B. thetaiotaomicron*. The colour scale reflects the abundance change of the most strongly changed protein annotated to that PUL. The fractions of proteins which are significantly up-/down-regulated are indicated in the heatmap. Only reliably measured (in at least 6 co-culture conditions) and strongly regulated (sum(abs(log_2_(fold-change))) > 4) PULs are shown. PUL annotation was obtained from (Ryan et al. 2024) which integrated data from (Martens, Chiang, and Gordon 2008; Martens et al. 2011; Briliūtė et al. 2019; E. D. Sonnenburg et al. 2010; Glowacki et al. 2020; Temple et al. 2017; Cartmell et al. 2017). (E) Number of proteins annotated to ‘Arginine biosynthesis’ which are up-/down-regulated in the presence of *C. aerofaciens*. (F) Number of proteins annotated to ‘Thiamine biosynthesis’ which are up-/down-regulated in the presence of *C. sporogenes*.

**Supplementary Figure 5.**
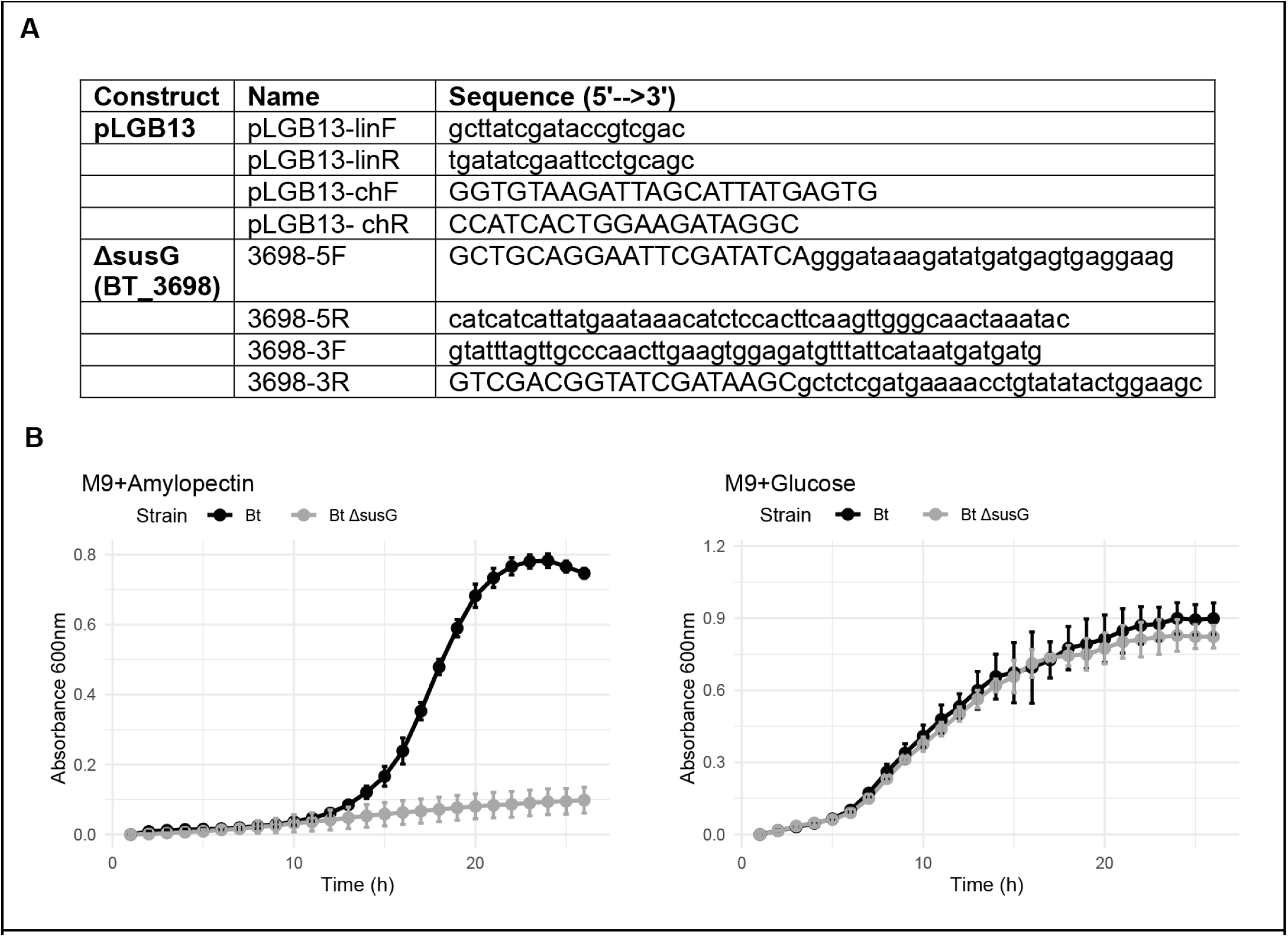
Starch utilisation mutant. (A) Table of primers used for constructing *B. thetaiotaomicron* susG mutant (Methods). (B) Growth curves in defined minimal medium containing amylopectin (left) and glucose (right) as the sole carbon source.

**Supplementary Figure 6.**
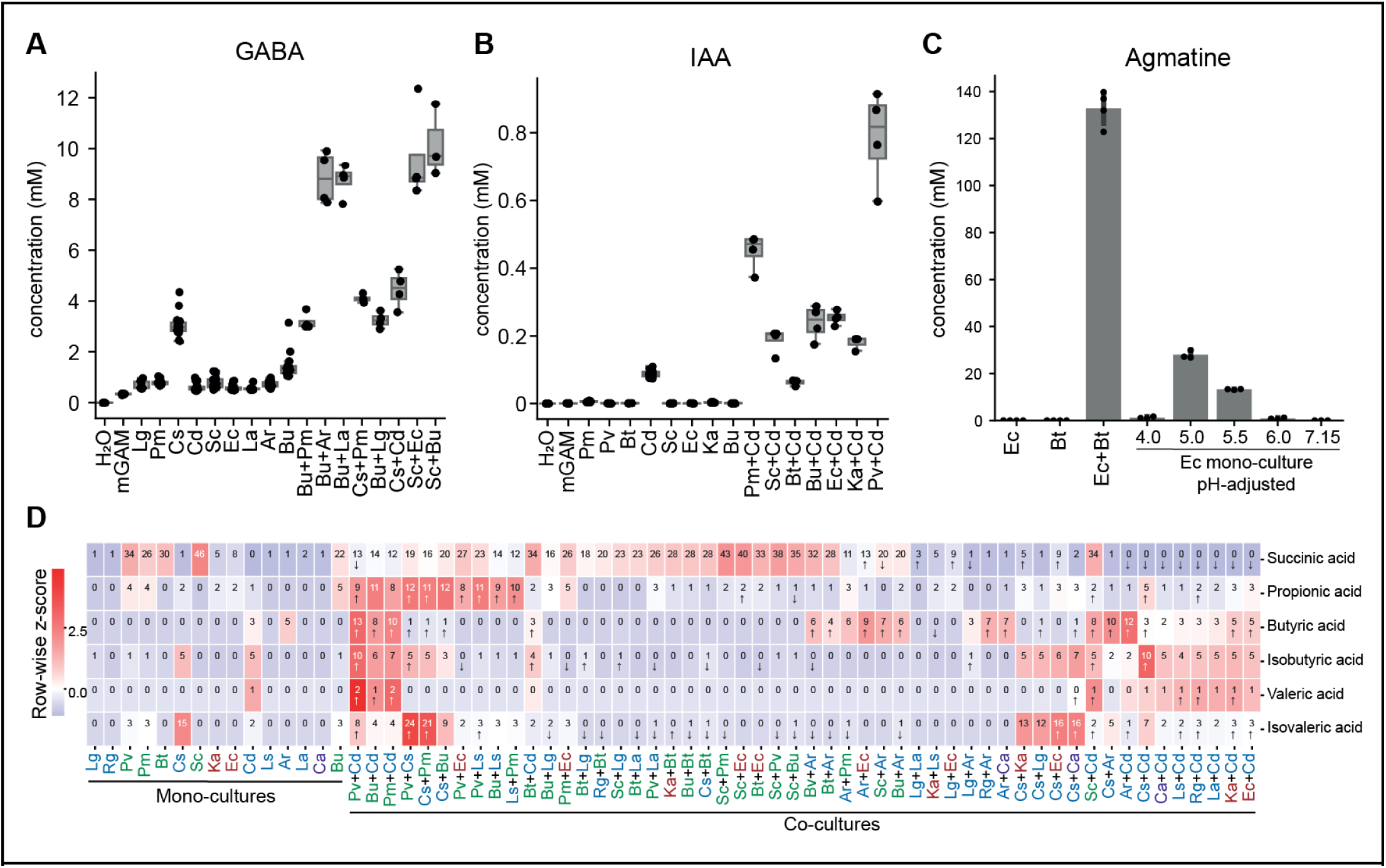
Additional metabolomics analyses. (A) γ-Aminobutyric acid concentrations from main metabolomics dataset for emergent co-cultures and relevant mono-culture conditions. (B) Indole-acetic acid concentrations from main metabolomics dataset for emergent co-cultures and relevant mono-culture conditions. (C) Agmatine concentration in supernatant in an independent small-scale experiment testing the effect of co-culturing and pH. Media pH was adjusted to the indicated value using hydrochloric acid. *E. coli* growth was substantially inhibited at pH 4, explaining the absence of agmatine. (D) Cross-species interactions shape distribution of fermentation end products. Heatmap colours indicate row-wise z-scores. Numbers in the heatmap indicate the concentration in mM. Arrows indicate significant deviation from the expected value in the indicated direction. Many Bacteroidetes produce large amounts of succinic and propionic acid and this is not additive in co-culture indicating competition for precursors. Several species can stimulate butyric acid production by *C. difficile* and *C. sporogenes*. Isobutyric and isovaleric acid production is suppressed in several co-cultures potentially indicating competition for the amino acid precursors.

**Supplementary Figure 7.**
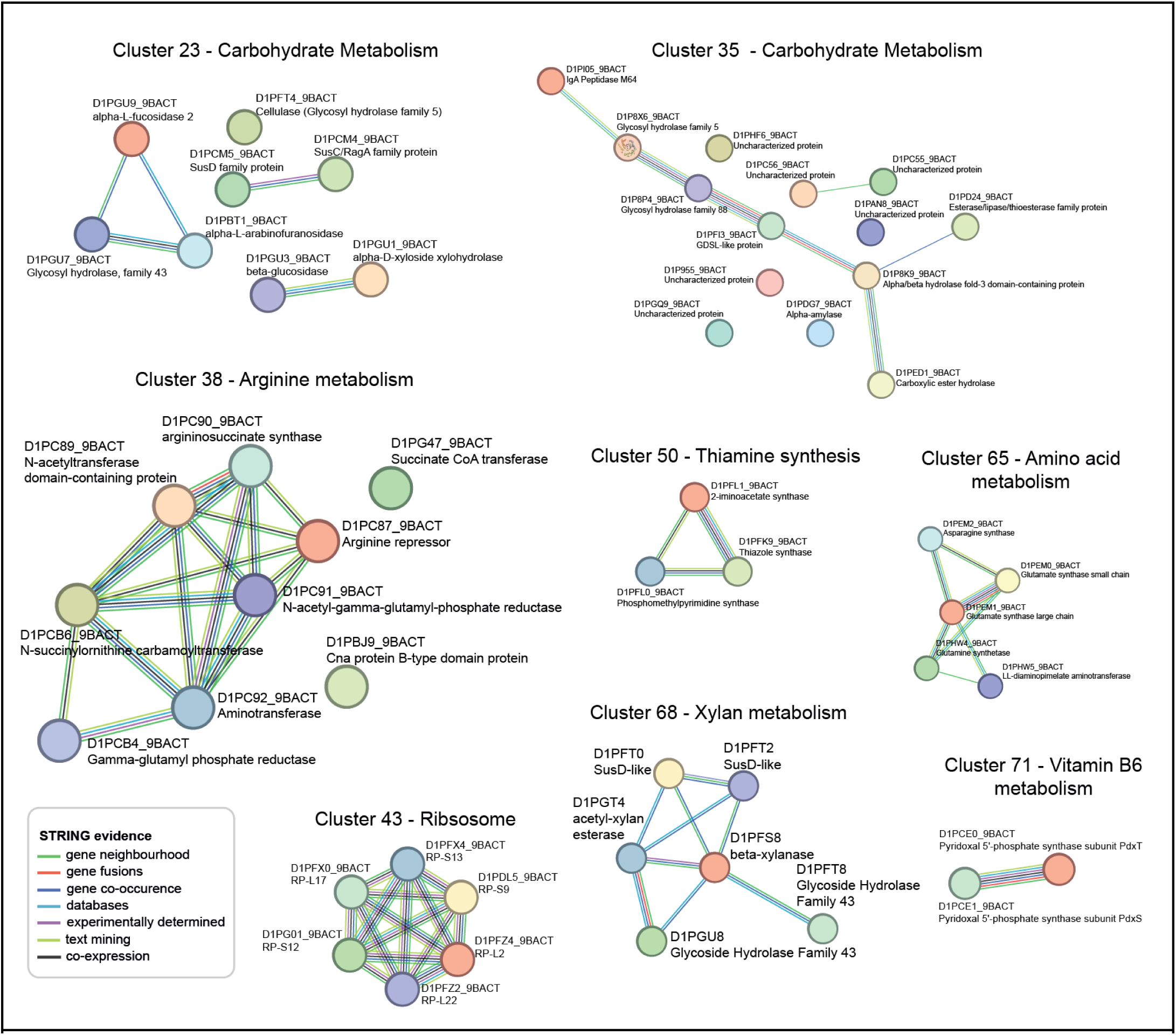
StringDB networks of *S. copri* clusters. Selected clusters of *S. copri* proteins (**Fig 6E**). Networks illustrating known connections between cluster members based on various types of data curated in StringDB.

## Notes

### Competing Interest Statement

The authors have declared no competing interest.

